# Directed networks and resting-state effective brain connectivity with state-space reconstruction using reservoir computing causality

**DOI:** 10.1101/2023.06.08.544175

**Authors:** Joan Falcó-Roget, Adrian I. Onicas, Felix Akwasi-Sarpong, Alessandro Crimi

## Abstract

Causality theory is a complex field involving philosophy, mathematics, and computer science. It relies on the temporal precedence of cause over a consequence or unidirectional propagation of changes. Despite these simple principles, normative modeling of causal relationships is conceptually and computationally challenging. Model-free approaches provide insights into large, complex, and chaotic networks, but suffer from false positive and false negative inferences caused by meaningless statistical and temporal correlations. Machine learning advancements have extended these data-driven methods to nonlinear systems, yet inherited similar drawbacks as linear approaches. Interestingly, newer proposals within this model-free paradigm reverse the temporal precedence using the internal structure of the driven variable to recover information from the driving one. Efficient machine learning models combined with these state space reconstruction methods automate part of the process, potentially reducing inductive biases during training and inference. However, their translation into neuroscience, especially neuroimaging, is limited due to complex interpretations and a lack of systematic analyses of the results. Here, we exploited these methods combining them with normative analyses to reconstruct chaotic relationships and networks emerging from neuroimaging data. We validated the proposed scores with a chaotic yet solved system and rebuilt brain networks both in synthetic and real scenarios. We compared our method and heuristics with well-established alternatives providing a comprehensive and transparent benchmark. We obtained higher accuracies and reduced false inferences compared to Granger causality in tasks with known ground truth. When tested to unravel directed influences in brain networks meaningful predictions were found to exist between nodes from the default mode network. The presented framework explores reservoir computing for causality detection, offering a conceptual detour from traditional premises and has the potential to provide theoretical guidance opening perspectives for studying cognition and neuropathologies.

**Author summary:** In sciences, reliable methods to distinguish causes from consequences are crucial. Despite some progress, researchers are often unsatisfied with the current understanding of causality modeling and its predictions. In neuroscience, causality detection requires imposing world models or assessing statistical utility to predict future values. These approaches, known as model-based and model-free, have advantages and drawbacks. A recent model-free approach augmented with artificial networks tries to autonomously explore the internal structure of the system, (i.e, the state space), to identify directed predictions from consequences to causes but not the other way around. This has not been extensively studied in large networks nor in the human brain, and systematic attempts to reveal its capabilities and inferences are lacking. Here, the proposal is expanded to large systems and further validated in chaotic systems, challenging neuronal simulations, and networks derived from real brain activity. Although the manuscript does not claim true causality, it presents new ideas in the context of current trends in data-driven causality theory. Directed networks encoding causality are hypothesized to contain more information than correlation-based relationships. Hence, despite its evident difficulties, causality detection methods can hold the key to new and more precise discoveries in brain health and disease.

## Introduction

Strikingly, the activity of certain regions of the (not only) human brain was found to be suppressed in task-induced engagement [1]. In addition, brain activity has been demonstrated at rest [2], first observed using positron emission tomography, and later in functional Magnetic Resonance Imaging (fMRI) studies [3]. This line of research further developed to define what is currently known as resting-state networks, such as the default mode network (DMN) [4]. The regions comprising DNM are symmetrically distributed across the association cortex [5]. Extensive and ongoing research is devoted to highlighting the role of this subset of intertwined regions in a variety of cognitive domains [6]. However, high-resolution fMRI studies addressing individual variability suggest the existence of multiple sub-network within the DNM [7].

To this end, understanding how patterns of coordinated activity across these populations of neurons are organized is central to the study of brain function in health and disease. Functional MRI is a widely used neuroimaging modality for mapping synchronous activity between brain regions, which typically relies on identifying bidirectional connections across brain regions showing similar oscillations across time. Yet, functional communication is hypothesized to take place unidirectionally, with the dynamics of one region influencing the occurrence of events in another region. This is indicative of information being transmitted casually. This phenomenon is defined as directed, or effective connectivity [8], and is *typically* recognized by the presence of a delay in the temporal correlation of the spontaneous signal oscillations of brain regions.

Untangling directed influences instead of simple correlations within the DMN shows great potential for further improving the overall comprehension of brain function in health, disease, and development. Effective connectivity within the DMN has received attention in the last decade, however, with little consensus on the directionality of the functional interactions. Noteworthy, these studies have primarily focused on small and nongranular representations of the network [9]. Recent research acknowledges stronger influences from posterior to anterior regions rather than the opposite [10, 11].

Crucially, causality itself and its implications to neuroscience are not and probably will never be exempt from controversy [12] due to difficulties both in the philosophical and practical [10] domains. Despite this, several techniques have been used throughout its history. In what follows, we provide a (necessarily) brief review that led us to propose the framework we study in this manuscript and its application.

### Autoregressive modeling of asymmetric influences

Granger influence modeling, which in some scenarios is equivalent to information-theoretic quantities [13], is a statistical concept that measures directed influences between at least two-time series [14]. In short, Granger Causality (GC) tests whether the past values of at least one time series help in predicting the future values of another time series. A cause (or a discrete set of causes) is a reliable and significant predictor of a consequence. In practice, the main component of GC is the estimate of autoregressor variables which are then further validated by F-statistics. These approaches are data-driven and do not require the addition of prior knowledge as opposed to structural and dynamic causal modeling [15, 16] or diffusive approaches [17]. We refer the interested reader to insightful reviews on these techniques [18, 19].

However, due to the potential confounding characteristics that each regressor may entail (see for example [18, 20, 21]), there are still ongoing disagreements regarding the definition of causal interaction between brain regions [22] using this framework. Some authors consider GC-based approaches as primarily being temporal association measures [23]. Within the same paradigm, interesting extensions are aimed to improve the prediction skill between nonlinear relationships by using deep layered networks [24–27]. Furthermore, to unmask spurious causalities, given by unseen variables or temporal correlations, a possible approach can be given by studying changes in the system given by observable perturbation for instance by using transcranial magnetic stimulation [28]. Unfortunately, these settings are difficult to achieve in all circumstances, both *in-vivo* and *in-silico*. We defer further arguments about this aspect and our proposed method in the Discussion.

It is also recognized that one limitation of GC originates from non-stationary time series [29]. Importantly, it has been suggested that stationarity should not be naively assumed in time series of brain activity [30]. This is supported by experimental findings using different neuroimaging techniques, from electrophysiology in high order mammal species [31], electro- and magnetoencephalography [32, 33], and fMRI [34, 35] in humans. These non-stationary patterns of activity have been further characterized with computational models of coupled dynamic systems. Inherent noise permits the brain to explore the surroundings of stable attractors [36, 37], potentially encoding functionally relevant resting-state networks [38] and transitions between them [39, 40]. Thus, the stationarity within resting-state networks is strongly challenged [35]. Nonetheless, having acknowledged these limitations, we still accept the fact that Granger influence modeling can give us plausible and valuable insights when modeling brain effective connectivity in controlled environments [41, 42].

### State space reconstruction allows for unidirectional mappings

Another line of research is committed to finding answers to the same challenges discussed above but focusing on the properties of the state space of the system at hand rather than the data itself. State Space Reconstruction (SSR) methods leverage Takens’ theorem [43] to find unidirectional mappings between two variables in the neighboring points of the system’s attractors (see pp. 6 in [44] for a very insightful summary). In brief, the theorem states that the state space of the dynamical system {*x_i_*}*_i_*_=1,2,3_*_,…_* is homeomorphic to a shadow manifold built from delayed time points of any of the system variables *M_xi_* ={*x_i_*(*t* + *kτ*)}*_k_*_=0,1,2,.._ In a dynamical system, Cross Convergent Mapping (CCM) [45] exploits the fact that a uniform mapping ”*∼*” can be found from the driven variable to the undriven one between their shadow manifolds *M_Xi_ ∼ M_Xj_* . In plain words, if using the aforementioned mapping *X* can be estimated using *Y* better than the other way around, then *X* can be said to CCM-cause *Y* . Importantly, this represents a paradigm shift regarding GC-based methods but at the same time poses a challenge for the comprehension of results derived from these approaches [45].

Although proven to disentangle causal effects in chaotic and highly nonlinear systems, CCM is not immune to pitfalls even producing incorrect results in fairly intuitive systems [46]. CCM-causality is parameter-dependent and requires asymptotic behavior concerning time series length. Moreover, it is not trivial to find the embedding dimension nor the particular lag *τ* at which the mapping is optimal. Lastly, to the extent of our knowledge, its performance in non-stationary systems is yet to be studied.

Several extensions of CCM aimed to solve these and other related issues have already been developed [44, 47] yet without overcoming the hyperparameter dependence. Of particular relevance is the case of the Reservoir Computing Causality (RCC) variation [48]. Despite building on the same conceptual framework, RCC takes advantage of the superior forecasting power of Reservoir Computing Networks (RCNs) to autonomously build the manifolds, find the mappings and reconstruct the target time series. Hence, the embedding dimension as well as the lag are automatically found during the training process. Despite this, a new set of hyperparameters needs to be adequately selected, potentially inducing biases during the process. Fortunately, compendiums already exist with the sole purpose of guiding the choice of reservoir architecture design while at the same time increasing the network’s robustness to noise-related performance decrease [48, 49]. Noteworthy, the former is particularly appealing when dealing with MRI data.

In the present work, we test the ability of RCC to discover directionality in noisy time series. We developed a simple yet systematic and unambiguous procedure to extract statistically significant asymmetry in dynamic systems (Fig. 1). In contrast to the original proposal [48], this allows us to automatically explore causal relationships in large networks with unknown delays. In what follows, we provide a concise introduction to RCNs, we present a normative account of RCC and how the information derived from it can be used to reconstruct directed networks. Later on, in the Results we show the performance of the whole workflow applied to synthetic yet realistic systems and resting neuroimaging data from the previously mentioned DMN. Finally, we discuss our findings and provide potential intriguing directions that can be followed from hereon.

**Fig 1.**
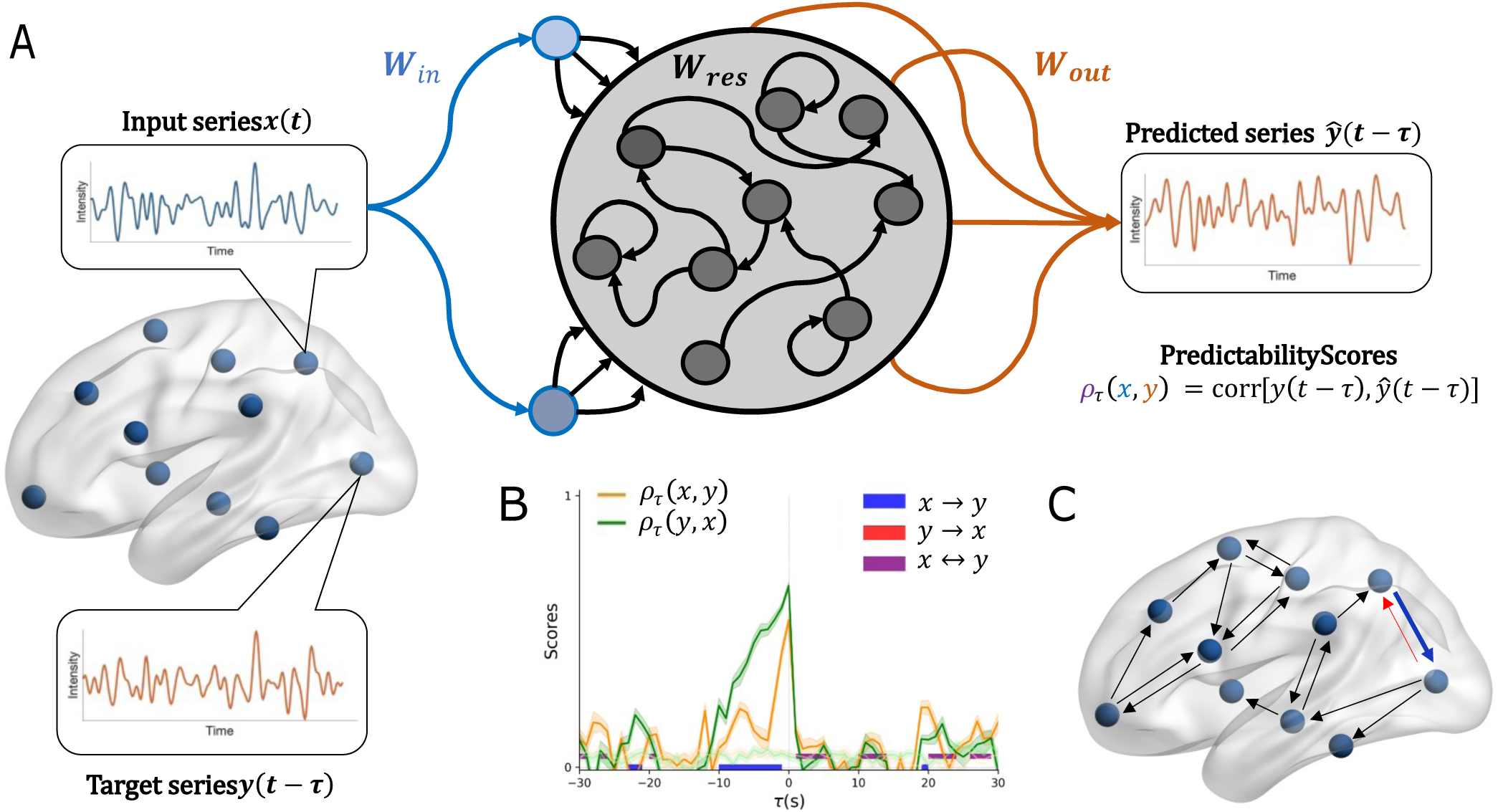
Reservoir Computing Causality framework for fMRI time series. **A** Temporal series extracted from two regions of interest (ROIs) are treated as input and target respectively. Once the reservoir is trained to predict the desired *τ* -lagged target, a predictability score is obtained. **B** The previous steps described in A are repeated for a sufficient number of negative and positive lags *τ* . The input and target series are then swapped and the same proceeding is applied to obtain the predictability scores. Several surrogates are generated to assess the significance of the obtained scores. **C** Between the two ROIs, a directed edge is assigned depending on the significant evidence in B. All these steps are then repeated across the desired nodes in the network to obtain a directed graph.

**Fig 2.**
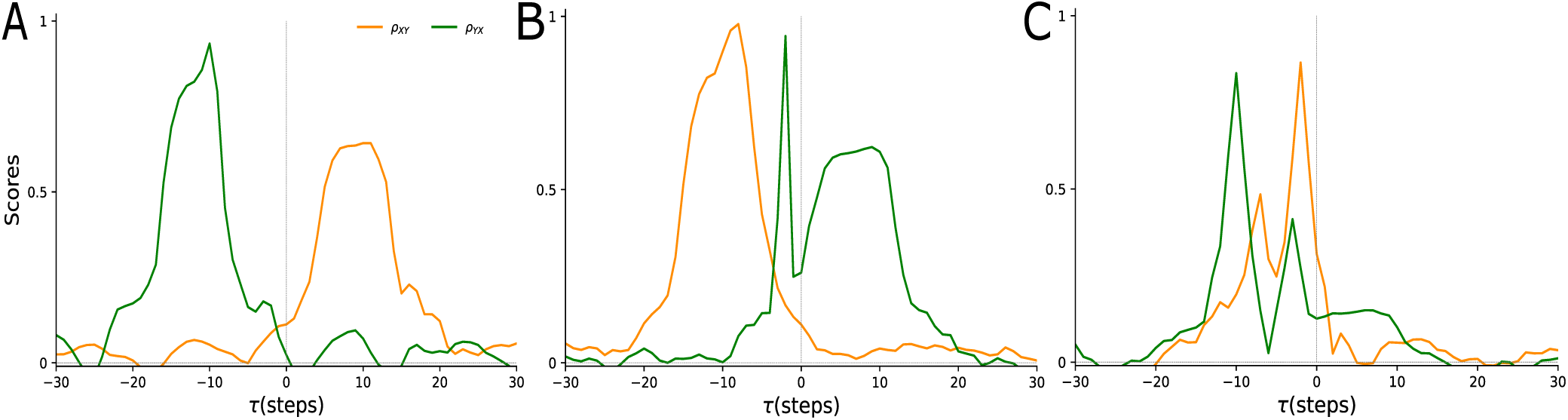
Reservoir computing causality for *noise-free* logistic mappings. The exact values of the couplings can be found in 3. **A** Simple unidirectional coupling *x → y* is present at *τ* = 10. **B** Bidirectional coupling *x* ⇄ *y* found at *τ* = 2 and *τ* = 8 respectively with different intensities. Recall how the evidence for the direction *x → y* is found at both positive and negative *τ* . **C** Double bidirectional coupling with different intensities. Recall how the small green peak would not be detected by the method presented in the main text. Yet, peak analysis is simply not meaningful in real data given the obvious deviation from the ideal cases presented here (see Fig. 3).

## Materials and Methods

### Reservoir Computing Networks (RCNs)

The basis of the method we propose relies on assessing how predictable delayed effects are between two distinct observations. Predictions are made with RCNs which, in general, consist of three building blocks. An *Input-to-node* module accepts the input series and feeds it to the *Node-to-Node* (or *reservoir*). Finally the *Node-to-output* module retrieves the information producing a desirable output (Fig. 1A). We used the newly developed PyRCN library written in Python [50] which provides easy-to-use workflows to build and test different RCNs.

At a given time step *t*, the *N_in_* units in the *Input-to-Node* connect an input sample *x_t_* to the reservoir via a set of sparse weights **W***_in_*,

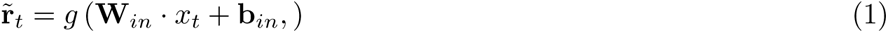

where *g*(*·*) is any non-linear function (including the identity) and **b***_in_* are a set of biases. This high-dimensional representation of the input features 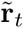 is fed to the whole set of *N_res_* units present in the main body of the reservoir.

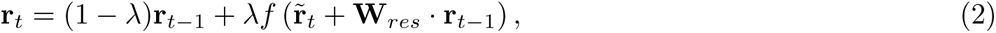

where, again, *f* is introducing a non-linearity and the leakage *λ ∈* (0, 1] controls the contribution of the previous reservoir states **R** = {**r***_n_*}*_n_*_=1*,…,t*_*_−_*_1_ to the current state *r_t_*. The prediction (i.e., the output of the RCN) of the *τ* -lagged target sample is a linear readout of the reservoir state expanded by a constant of 1:

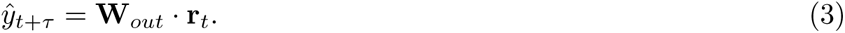

Only the output weights **W***_out_*, containing a set of biases, are then trained using regularized linear regression to adequately match the desired target series [50]. Finally, the predictability of the entire *τ* -lagged target series **y***_t_*_+*τ*_ given the predictor **x***_t_*is assessed by the Pearson correlation between the target and the predicted series

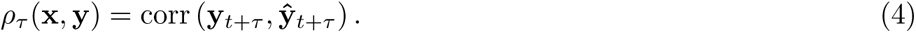

Shortly, the main idea behind reservoir-like computing is that a given input pushes the reservoir to specific locations in a high-dimensional manifold [51, 52]; the output weights are then optimized to retrieve information from the nearby regions. Were the input to move the reservoir away to other points, the output weights would not be able to recover meaningful information hence completely missing the prediction. As simple as this paradigm seem, there is further evidence that RCNs supersede state-of-the-art deep learning-based models for temporal series prediction even on the verge of chaos [53]. Interestingly, richer approaches aim to train the reservoir connections themselves to increase the have been proven to be useful to understand the dynamical properties of cortical networks exist (see [54] for an extensive review).

Briefly, the concept behind reservoir computing is to process and transform input data using a reservoir of nonlinear dynamical nodes, analogous to how complex biological systems, such as the brain, process and transform data. It does not, however, directly represent any one bodily system. Some academics have hypothesized that the temporal dynamics of the reservoir may resemble the neural activity in particular brain areas [55]. For instance, the distributed and heterogeneous connectivity present in biological systems can be modeled using the random links between elements in the reservoir. In this direction, an interesting proposal has directly mapped the brain white-matter connectome to the reservoir [56].

### Reservoir Computing Causality (RCC)

Phenomenologically, evidence for a delayed unidirectional coupling between two observations **x** *→* **y** exists if the maximum of the predictability *ρ_τ_* (**y**, **x**) occurs at negative lags *τ <* 0 while the mirrored score *ρ_τ_* (**x**, **y**) peaks at positive lags *τ >* 0 [47, 48]. Despite seeming counterintuitive this would mean that current information of **y** is encoded in previous observations of **x** and, consequently, that current information of the cause **x** explains future observations of the consequence **y** (see [45] for a comprehensive explanation). In some systems where the interactions are bidirectional, the rationale differs in that both predictability scores peak at negative lags *τ <* 0 being the height of the peaks informative of the coupling strength. However, the existence of this bidirectionality does not necessarily invalidate the previous phenomena. Instantaneous relationships violate the so-called *causality principle* which states that the cause precedes the effect, hence there is no point in inferring causality without an explicit delay (i.e., *τ* = 0).

In what follows, we present a method to systematically evaluate directionality in large networks. Our main hypothesis comes from the fact that predictability scores do not necessarily reach a clear and distinct maximum anywhere. This is especially relevant in BOLD time series where noise plays a determinant role and the length is limited due to acquisition challenges (e.g., expensive). We thoroughly consider the minimum requirements that need to be met for a given result to indicate the presence or absence of both directionality and bidirectionality between nodes in a network.

Recall that the evidence for directionality is self-contained in the difference between mirrored predictability scores. Following McCracken and Weigel [46], we define a Δ-score,

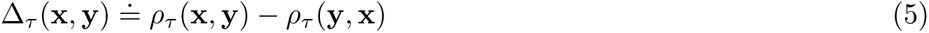

where Table 1 summarizes all possible situations discussed previously.

**Table 1.**
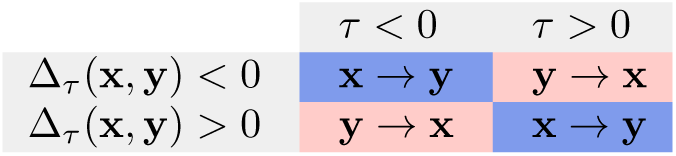
Summary of the interpretation of the Δ-score proposed in [46] and further extended here self-containing the evidence for directionality in the RCC framework. Nonetheless, its interpretation is far from intuitive.

### Reconstruction of directed networks

Let us now consider an example of **x** *→* **y** tested in the ***negative*** *τ* regime. The minimum things required to consider directionality are the following:

1. a statistically significant predictability of the **x** series from **y** with respect to its surrogate relationship and,
2. a statistically significant evidence for negative Δ-scores with respect to the surrogate score; specifically, the Δ-score computed using the predictability scores corresponding to each of the two surrogates **x***_s_* and **y***_s_*.

Based on the previous example and the summary present in Table 1 it is possible to extrapolate these arguments to the general case (see S1 Appendix). Therefore, we define two lag-dependent *δ_τ_* -scores encapsulating directionality as follows:

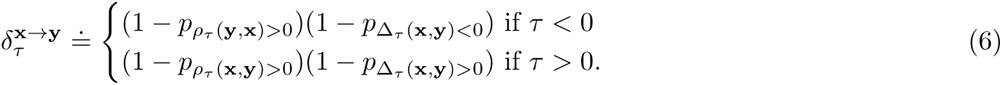

Similarly, the score for the opposite direction can be defined as:

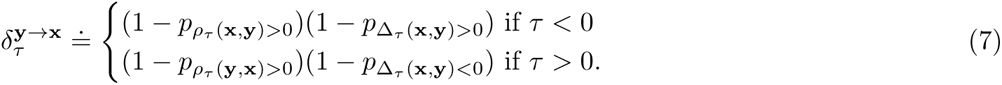

Note how the first bracket takes point into account 1) while the second bracket encapsulates point 2). Importantly, these scores are not symmetric nor antisymmetric, meaning that directionality is not easily interchangeable and is solely dependent on the data as well as the fitness of the trained RCN 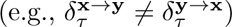. Furthermore, we can also extract bidirectionality by defining

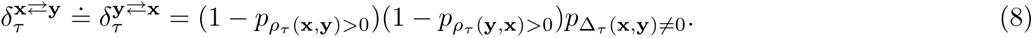

where *p_H_*_1_ is the *p*-value after testing the alternative hypothesis *H*_1_.

To assess statistical significance and compute the necessary *p*-values, along every input-target pair we generate surrogate time series of the target series to train the RCNs [57, 58]. From a given time series, we denote its surrogate with a subscript; hence, for a given series **x**, the corresponding surrogate is **x***_s_*. For every real input-output pair, we trained 20 different RCNs and generated 100 surrogates for each direction tested (i.e., **x** *→* **y***_s_* 100 times and vice-versa). For each input-surrogate pair, only 1 reservoir was trained: as a result, the null distribution for a given pair of time series consisted of 100 samples for each separate direction (Fig. 1B). Finally, given the unequal sample size, we used Welch’s t-test (i.e., without assuming equal variances) to compute the 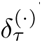 scores presented above.

As a final step in the process, the elements of the adjacency matrices **A***_τ_* are built using the aforementioned scores. Defining the *x*-th and *y*-th nodes of the network, whose time series are **x** and **y** respectively, each entry is obtained as

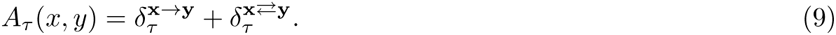

Although unlikely, there might be a case where there is both evidence for mutual influence and a specific direction. However, simultaneous evidence for both directions is not possible independently. In any case, the bidirectional component can be dropped if only the asymmetric part is informative; that is,

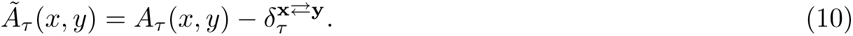

Retaining the symmetric component of the network might be relevant in the case of resting-state brain networks, where the aim is not only to reconstruct the directed part but also several co-activation between spatially separated brain regions. Yet, in both scenarios, it follows from Eqs. (6) and (7) that the reconstructed networks are directed but not signed (Fig. 1C).

### Ground truth validation

The previously defined scores are based upon statistical hypotheses and corresponding *p*-values. As a result, they are normalized between 0 and 1, with the latter indicating a clear (bi)directionality. Therefore, it is straightforward to extract binary networks based on statistical evidence. We obtained the thresholds to consider the scores in Eqs. (6), (7) and (8) as significant by replacing the *p*-values for their significant levels. We also corrected for multiple hypotheses testing: 3 for Eqs. (6) and (7) and 2 for Eq. (8).

However, a more thorough analysis of the results comes from treating the entries of reconstructed adjacency matrices in Eq. (10) as probabilities of classifying a given edge as existing or non-existing. Within this framework, we computed Positive Predictive Values (PPVs), Negative Predictive Values (NPVs), sensitivity, specificity, and finally, Area Under the Receiving Operating Curve (AUROC) scores. As it is common in machine learning, for each time series we used 70% of the length to train the RCNs and then used the other 30% to test them, that is, all scores computed were obtained not on the training time points but rather on the testing time points.

### Resting-state fMRI data

The Population Imaging of Psychology 2 (PIOP2) dataset [59] consists of 8 min of rs-fMRI (240 timepoints at 2-second TR) from 226 subjects (mean age = 21.95 *±* 1.78 years, 57% females, 89% right-handed). Data were preprocessed in the volumetric space using the default fMRIprep pipeline [60], a Nipype [61] based tool focused on transparent and reproducible preprocessing for neuroimaging data. Each functional image was denoised using AFNI’s 3dTproject, including the following [62]: (1) 6 motion parameters (3 rotations and translations in x y, and z directions), (2) global, white matter, and cerebrospinal fluid signal, (4) derivatives, squares and squared derivatives of each noise regressor, (5) framewise displacement motion spikes (see below), (6) linear and polynomial trends, and (7) band-pass filtering to exclude frequencies outside of the range 0.01-0.15 Hz and improve the signal-to-noise ratio.

Head motion was estimated using the framewise displacement (FD) relative root mean squared (RMS), which was calculated using FSL MCFLIRT [63]. Motion spikes were defined as FD threshold *≥* 0.25 mm and were included in the model as dummy variables (i.e., spike regression; [62]). Scans (*n* = 1) were excluded for gross motion if they had an average FD RMS *≥* 0.20 mm or *>* 38 (15%) volumes with FD RMS *≥* 0.25 mm [62]. Two scans were discarded due to signal dropout. Cortical regions of interest (ROIs) were defined based on an atlas of the human brain’s functional connectivity architecture, derived from 1489 participants using gradient-weighted Markov Random Field models. The resulting atlas consists of a 100-ROI parcellation divided into 7 intrinsic functional networks [64]. For the current study, the default mode network (DMN) comprising *n*=24 nodes was selected as a priori network of interest, constructed by extracting average time series within each node to study their effective relationships. Importantly, this represents a larger attempt to comprehend directed connectivity as opposed to earlier studies [9–11, 65–68].

### Data and code availability

All the material necessary to reproduce this study is openly available in the form of an Open Science Framework repository [69].

## Results

Despite more than 20 years since its original proposals [51, 52], RCNs have been rather eclipsed by other machine learning frameworks such as deep learning. An interesting line of research attempts to combine both by leveraging the efficiency of RCNs and the representational power of deep neural networks [70–72]. Given its easy implementation in PyRCN, we tried several combinations of them but, in all our experiments, we found no improvement in prediction accuracy and, in some cases, even returned non-convergence warnings. To this end, and keeping in mind that SSR methods rely on mappings in the embedding space, we designed a simple yet robust RCN with a rather small number of neurons. When training these networks with surrogate data, in some cases, invalid convergence values were found. This was expected since surrogate time series should, in some sense, be unpredictable from the original inputs. The specific parameters for all the networks trained can be found in Table 2. As already stated earlier, RCNs were trained on 70% of the length and tested on the remaining 30%. Small variations in this percentage split did not produce differences, provided that a sufficient number of training samples and, in fewer amounts, testing samples were available (i.e., 60-80%).

**Table 2.** Summary of the parameters chosen to train the Reservoir Computing Networks (RCNs) in this work. For the two different blocks, *NA* stands for Not Applicable, and *null* indicates that the value was left empty to be chosen by the implemented random sampler. For further details on the meaning of each one of these parameters we refer the reader to the original publication of the package [50] and documentation *here*.

| Object | Input-to-Node | Node-to-Node |
| --- | --- | --- |
| **Units (#)** | 50 | 50 |
| **Sparsity** | 1 | 1 |
| **Activation** | logistic | tanh |
| **Scaling** | 1 | NA |
| **Shift** | 0 | NA |
| **Bias scaling** | 1 | NA |
| **Bias shift** | 0 | NA |
| **Random seed** | *null* | *null* |
| **Spectral radius** | NA | 1 |
| **Leakage ($\lambda$)** | NA | 1 |
| **Bidirectional** | NA | *false* |

### RCC in corrupted logistic maps between two nodes

SSR methods often use a logistic coupling between two or more variables to test the validity of the method before showing applications to real data. These dynamical systems are known to exhibit chaotic behavior for certain sets of initial values making it particularly challenging for causal inference methods. Similarly to the original proposal in [48], but also previous works [44, 45, 47], we test the proposed method with a system of two coupled logistic series. Following the established trend, we generated the following two-node logistic network:

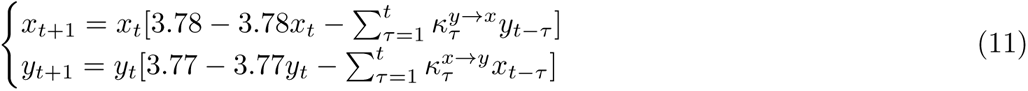

where the coupling constants 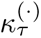 encode the directionality of the causal connections. To further illustrate the *intricate* nature of Reservoir Computing Causality (RCC) and other SSR methods in general we simulated three different noise-free logistic systems. Specifics of these simulations are in Table 3. The RCNs correctly identify the peaks where the predictability scores are maximum, hence disentangling causality from the generated time series. However, from these examples, it is clear that in high-dimensional systems of unknown directionality, a phenomenological approach is not feasible given the complex pattern expected in real scenarios.

**Table 3.** Numerical values for the coupling constants used to simulate the *noise-free* logistic two-node networks in Eq. 11. RCC reconstructed causalities are shown in Fig. 2.

| Fig. 2 | A | B |  | C |  |  |  |
| --- | --- | --- | --- | --- | --- | --- | --- |
| $\tau$ | 10 | 2 | 8 | 2 | 3 | 7 | 10 |
| $\kappa_{\tau}^{x \rightarrow y}$ | 0.8 | *None* | 0.02 | *None* | 0.01 | *None* | 0.02 |
| $\kappa_{\tau}^{y \rightarrow x}$ | *None* | 0.82 | *None* | 0.04 | *None* | 0.02 | *None* |

Importantly, we generate series resembling real data found in fMRI datasets. In general, fMRI data is relatively short in length and substantially noisy. For the example shown in the main text, the couplings are restricted to the following: 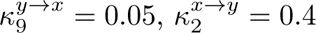 and zero for the rest. Hence, it follows that a double relationship exists at different delays. From this specific configuration, as stated in the main text, we generated 20 simulations that were treated as subjects, each one of them with different initial conditions. To corrupt the time series and test the method in *realistic* conditions, we only generated 250 points and corrupted the series with white noise as follows:

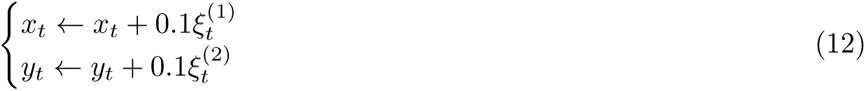

where *ξ*^(1)^, *ξ*^(2)^ are vectors containing random numbers drawn from a standard normal distribution.

Despite corrupting the time series, the RCNs correctly identify the causal connection if the delay is short (Fig. 3). However, the short length as well as the noise cause the predictability to fall significantly for larger coupling delays. As a result, in some subjects, directionality is completely missed (Fig. 3A right) while in others the identification is successful (Fig. 3A left).

**Fig 3.**
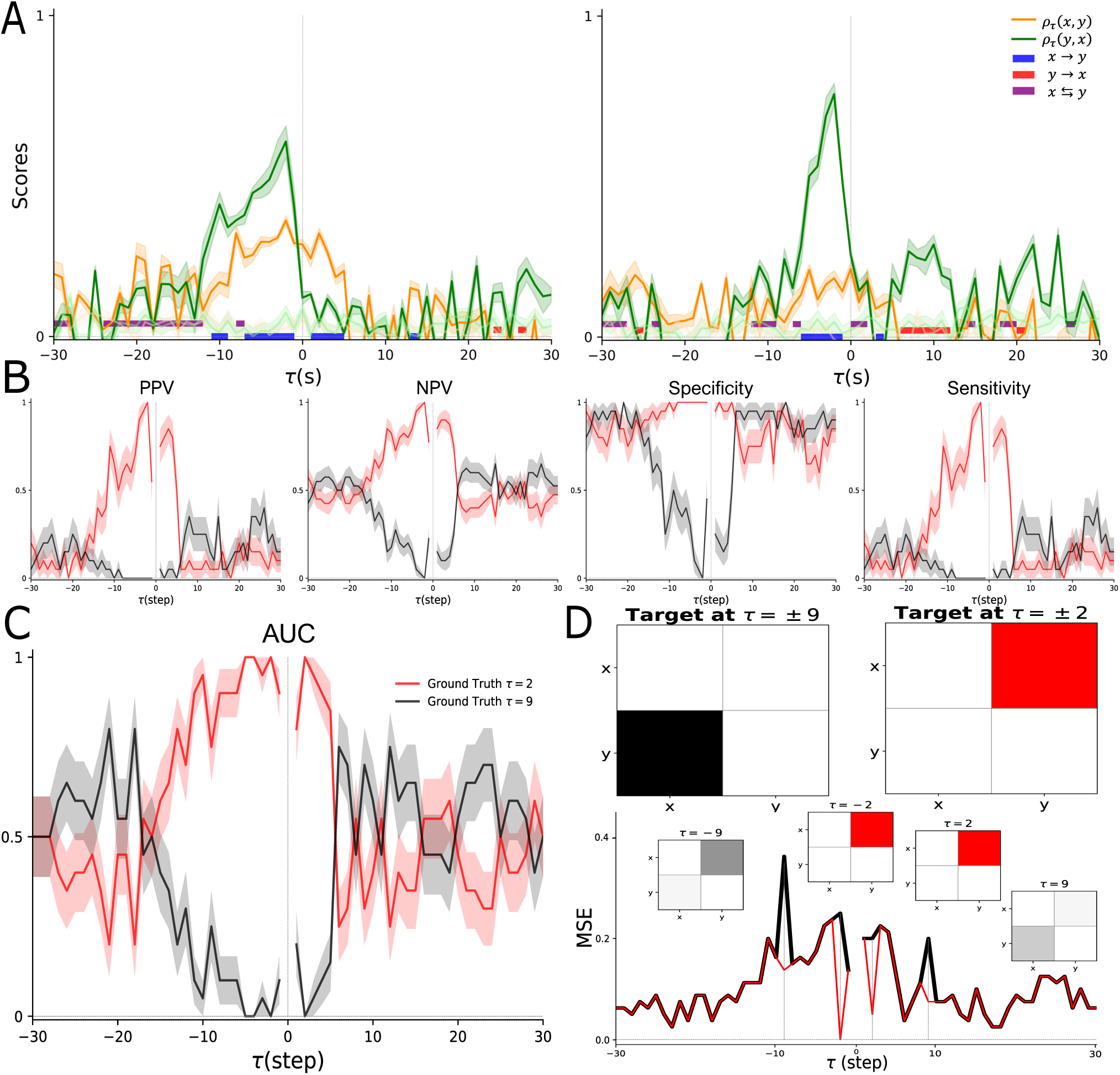
Reconstruction of the corrupted logistic 2-node network. All solid and shaded curves indicate results from the population analysis [mean*±*SEM]. **A** Predictability scores for two example networks. Transparent curves indicate the predictabilities of the corresponding surrogates. The colored bars below the curves mark significant evidence for all possible directionalities as defined in the main text. Note that the directionality from **B** Evaluation metrics as a function of the tested lag. For each *τ*, the reconstructed networks were compared separately with the two distinct ground truth networks generated. Each curve represents the score when comparing the reconstruction with the same ground truth across all tested lags. **C** Area Under the Curve (AUC) score for the task of correctly identifying the two distinct causal relationships. Note how one case mirrors the other meaning that the method correctly identifies one direction but fails to detect the opposite. Each curve represents the score when comparing the reconstruction with the same ground truth across all tested lags. **D** Visualization of the reconstructed networks obtained at all meaningful lags *τ* . At the bottom the Mean Squared Error between the reconstructed network and the ground truth which, besides the 4 targets shown at the top, consists of a matrix full of zeros. Note how directionality is not ubiquitously reconstructed in all lags hence causing the MSE to stay close to zero.

To further assess the predictive power of our approach, we divide the system in Eq. (12) in two sub-networks acting as ground truth: the networks containing 1) the short delay coupling and 2) the long delay couplings. In case 1) all the scores reach the maximum at the correct lag only in the case where the ground truth captures the short delay coupling (Fig. 3B red curves). Opposite to this, in case 2) the scores do not correctly reach high values at the desired lag *τ* = *−*9 but instead, decrease (Fig. 3 black curves). Moreover, the scores comparing the ground truth at short lags increase when the causal effect is not present. That is, the red curves increase at *τ* = *−*9 when they should decrease given that the correct ground truth corresponds to the network with the opposite directionality. In line with these scores, the method robustly identifies directionality across different decision thresholds but only when the causal effect is shortly lagged (Fig. 3C). In summary, the Mean Squared Error (MSE) across different lags between the reconstructions and the actual ground truth (i.e., containing only the connections in the correct delays) we find that directionality is correctly reconstructed only in the short lag case. Interestingly and importantly, positive lags appear to have similar predictive power as negative lags provided that the tested lag is short enough.

### RCC in realistic functional MRI simulations

We used a popular and rigorously tested dataset containing *realistic* fMRI simulations [73]. Netsim is a set of simulations designed with the specific purpose of testing numerical methods in computational neuroscience to unravel causality [18]. These simulations are also accompanied by the directed networks, of different sizes, upon which series for the different nodes were evolved in time. The original set of simulations contained 28 different simulated fMRI networks, each of them with specific underlying properties such as the Time of Repetition (TR) or the duration of the series. We studied a subset of those that contained the most representative variability to test both the reliability of our method as well as RCC within a neuroimaging domain (see Table 4). Finally, we compared the reconstructed networks using Eq. (10), containing only the asymmetric contributions, to the ground truth available *here*.

**Table 4.** Properties of the simulations used from the Netsim dataset. Specifications of the simulations chosen to test the Reservoir Computing Causality analysis method. TR stands for repetition time, an important parameter to consider when using functional fMRI time series. For further details, we refer the reader to the original publication [73].

| Simulation | Nodes (#) | Length (min) | Length (steps) | TR (s) | Noise (%) | Other |
| --- | --- | --- | --- | --- | --- | --- |
| 1 | 5 | 10 | 200 | 3.00 | 1.0 | neural lag=50 ms |
| 7 | 5 | 250 | 5000 | 3.00 | 1.0 | neural lag=50 ms |
| 15 | 5 | 10 | 200 | 3.00 | 0.1 | stronger connections |
| 19 | 5 | 10 | 2400 | 0.25 | 0.1 | neural lag=100 ms |
| 28 | 5 | 5 | 100 | 3.00 | 0.1 | neural lag=50 ms |

Although lagged-based methods are known to be challenging for these simulations, they are rather assertive when constrained to 1st-neighbor interactions [41]. To this end, and to avoid further discussions regarding the true nature of indirect connections [21], we also constrained our method to 1st-neighbors. In practical terms, this meant reconstructing edges in the networks that were known to exist *a priory*. For example, if a particular connection 1 *→* 2 existed in the real network, we computed scores in both directions and reconstruct both entries in Eq. (10). Alternatively, one may compute the RCC for every pair of nodes and mask the reconstructed networks by multiplying them by the binarized ground truth.

Although some SSR methods require a monotonous increase in the metrics as a function of the time series length [45, 46], this was not shown nor proven in the original proposal [48]. Nonetheless, we studied the dependence of the classification metrics on the length of the time series. For each time series, we used different percentages of the overall length and performed RCC, and reconstructed the networks. Noteworthy, the train-test splits remained in the 70-30% specified earlier. We found no clear dependence between the accuracy and the length of the time series for short lags (Fig. 4; see also S1 Figure). For small lags, we found that RCC was capable of achieving *better-than-chance* performance. Note that AUCs greater than 0.5 express the ability of the method to correctly classify a directed connection as existing or non-existing. For a more detailed analysis, please refer to the following sections.

**Fig 4.**
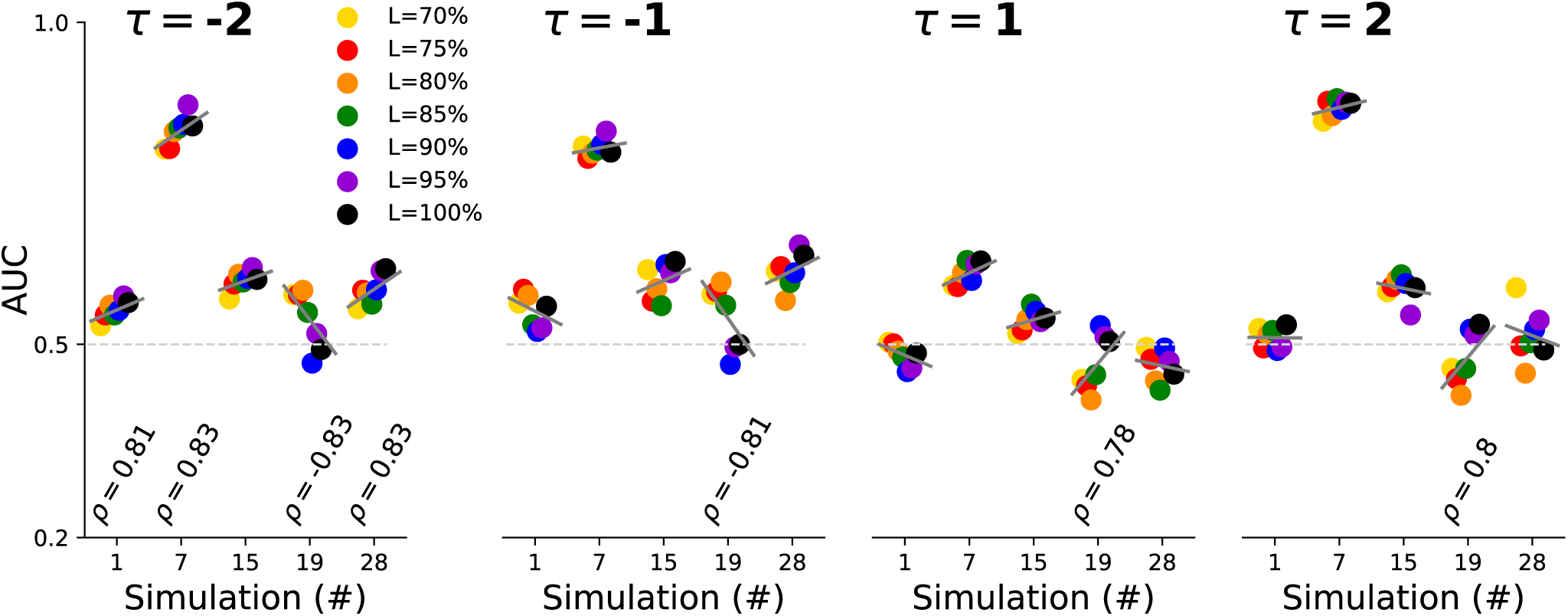
RCC’s Convergence of the classification metrics as a function of the time series length. Area Under the receiver operating Characteristic (AUC) for each simulation and the first 4 selected lags *τ* for each one of the simulations tested in this work. Short gray lines depict linear fits between AUCs and time series length. Numbers show the values of the statistically significant correlation coefficients found (*p <* 0.05, exact test). Error bars are not shown for the sole purpose of clarity.

### Improved directionality inference with RCC in realistic functional MRI simulations

We compared our analysis method with the result of Granger causality (GC), a well-established data-based method to infer directionality in stationary time series^1^. GC aims at unraveling directionality through an autoregressive linear model and a series of F-tests to assign a significance *p*-value for each edge [14, 29]. More precisely, let *x_t_* and *y_t_*be two extracted time series, in our case, from two different simulated nodes such that a prediction *ŷ_t_* is fitted using the following variate autoregressive model,

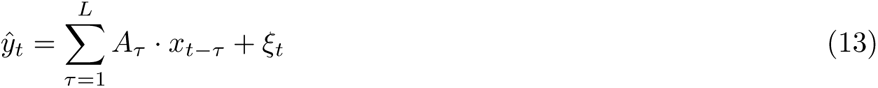

where *ξ_t_* introduces white Gaussian noise. The time series *x_t_* is said to Granger cause *y_t_* if *at least* one of the elements *A_τ_* for *τ* = 1*, …, L* is, in absolute value, significantly larger than zero. Given the autoregressive nature of GC, and to ensure a fair comparison with RCC, we simplified the canonical multivariate system [20] and included only two series per fitted model, as seen in Eq. (13). Furthermore, unlike GC, RCC tests for cause-consequence in a *one-lag-at-a-time* fashion, where directionality is accepted or rejected at specific time lags *τ* . To be consistent with this approach, we evaluated GC in a similar manner: we fitted *L* independent models. For each *τ* = 1*, …, L*, we evaluated the significance of the obtained coefficients *A_τ_* by defining a classification score based on the opposite of the measured *p*-value (i.e., 1 *− p_τ_*). Finally, we also constrained our analysis to 1st-neighbors in the same exact way as explained previously in the case of RCC. However, GC can not obtain effective connections for positive lags, hence making completely impossible a comparison between the two methods in the *τ >* 0 domain.

As expected, GC’s ability to detect directionality was independent of the length of the time series used (see S2 Figure); therefore, being the accuracy of RCC also independent of the length, we quantified the improvement in the classification metrics of our method with respect to GC taking into account only the results obtained with a length of 100% (i.e., using the whole length of the time series). However, we provide the results for all the different lengths in the supplements.

We computed the significance of the relative increase in PPVs, NPVs, and AUCs of RCC with respect to GC (Fig. 5). In all but one simulation, the combination of RCC and the analysis method proposed achieved higher scores than GC. Simulation 19 was the only one where RCC was not superior to GC, yet the performance was comparable.

**Fig 5.**
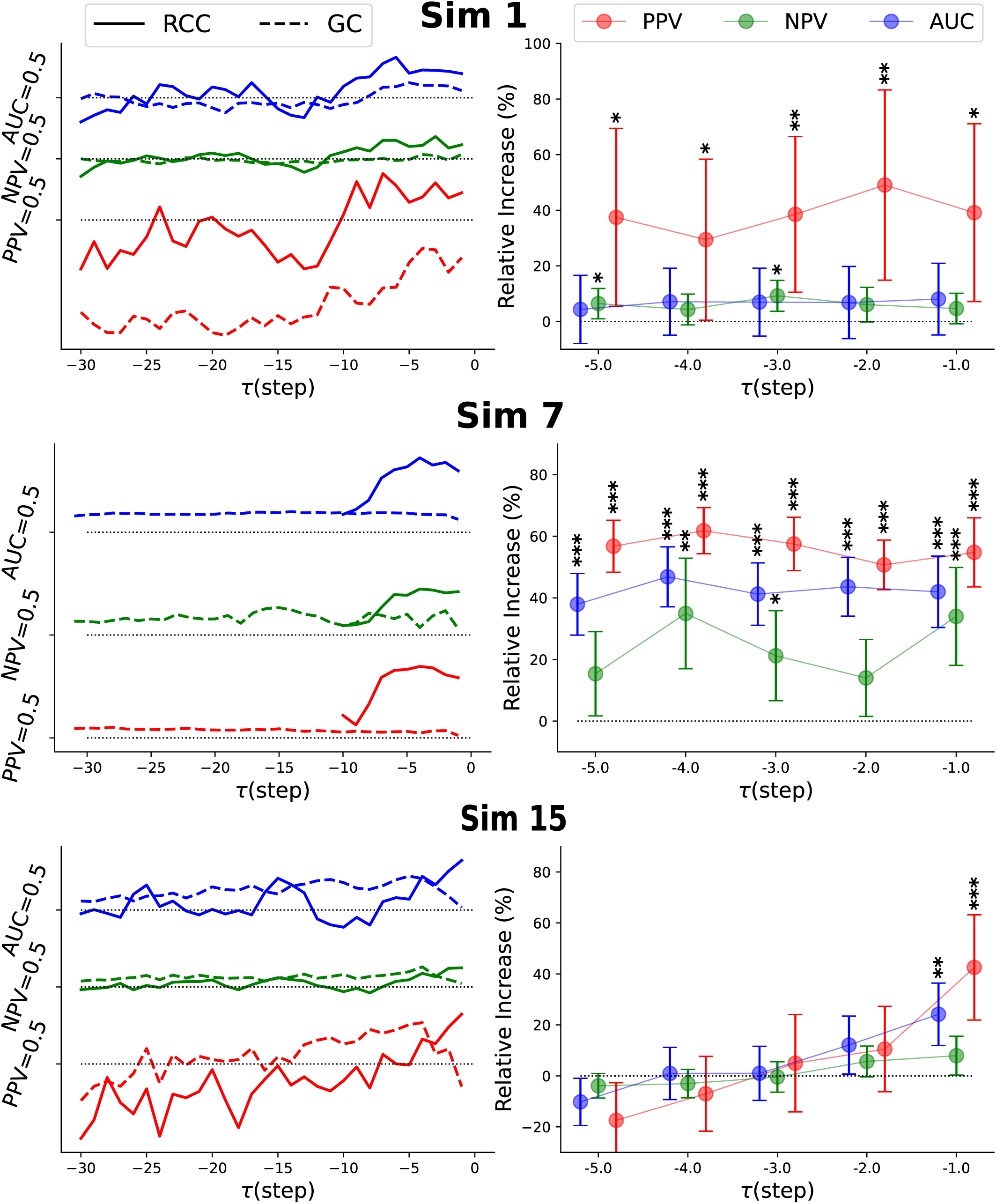
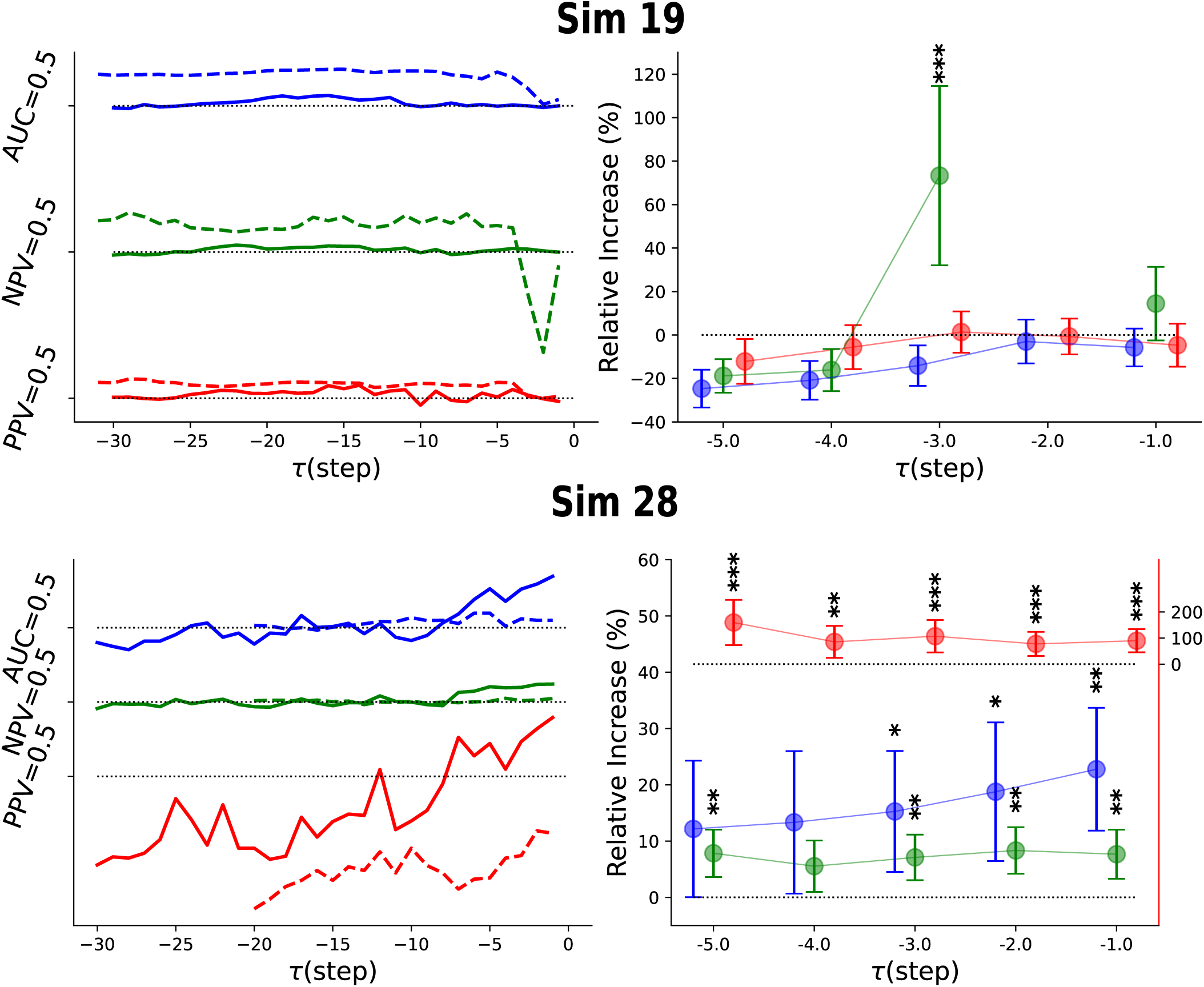
Comparison of RCC and GC in the Netsim dataset. LEFT Mean metrics of RCC and GC for each simulation tested for all negative lags. Positive Predictive Values (PPVs) are consistently and almost always significantly better for RCC. Importantly, the improvement of the combination of RCC and the proposed analysis method vanishes for sufficiently long lags in a similar manner as GC. In all plots, the dashed black lines indicate the *better-than-chance* threshold. RIGHT Relative increase of metrics from RCC with respect to scores obtained with GC for the first 5 lags accompanied with the corresponding significance analysis. Statistical significance was assessed using one-sided Welch’s t-tests given the hypothesized improvement of RCC in directionality detection (* for *p <* 0.05, ** for *p <* 0.01, and *** for *p <* 0.001).

Interestingly, the increase seemed to be *τ* -dependent. For large lags, RCC remained identical to the *better-than-chance* threshold indicating that the reservoirs did not induce recurrent trends nor were overfitting the synthetic data. Furthermore, the method’s ability to detect true positives was consistently higher than that of true negatives. Particularly noteworthy was the case of simulation 7 where the overall improvement was beyond statistical doubt. This was probably due to longer simulation times.

### Causality encoded in negative and positive lags

As stated previously, RCC finds directionality evidence both in negative and positive lags. In the previous section, we have shown how RCC compares to GC only in the negative lags range. Here we set out to study how the ability to detect directed connections using RCC differed between negative and positive lags (Fig. 6). We hypothesized that metrics would be higher in the negative lag regime given the evidence from the corrupted logistic map (see Fig. 2 and Fig. 3). In some simulations and small *τ* RCC performed significantly better in the negative regime. However, the overall improvement in the metrics was less prominent, rather erratic, and strongly dependent on the simulation number.

**Fig 6.**
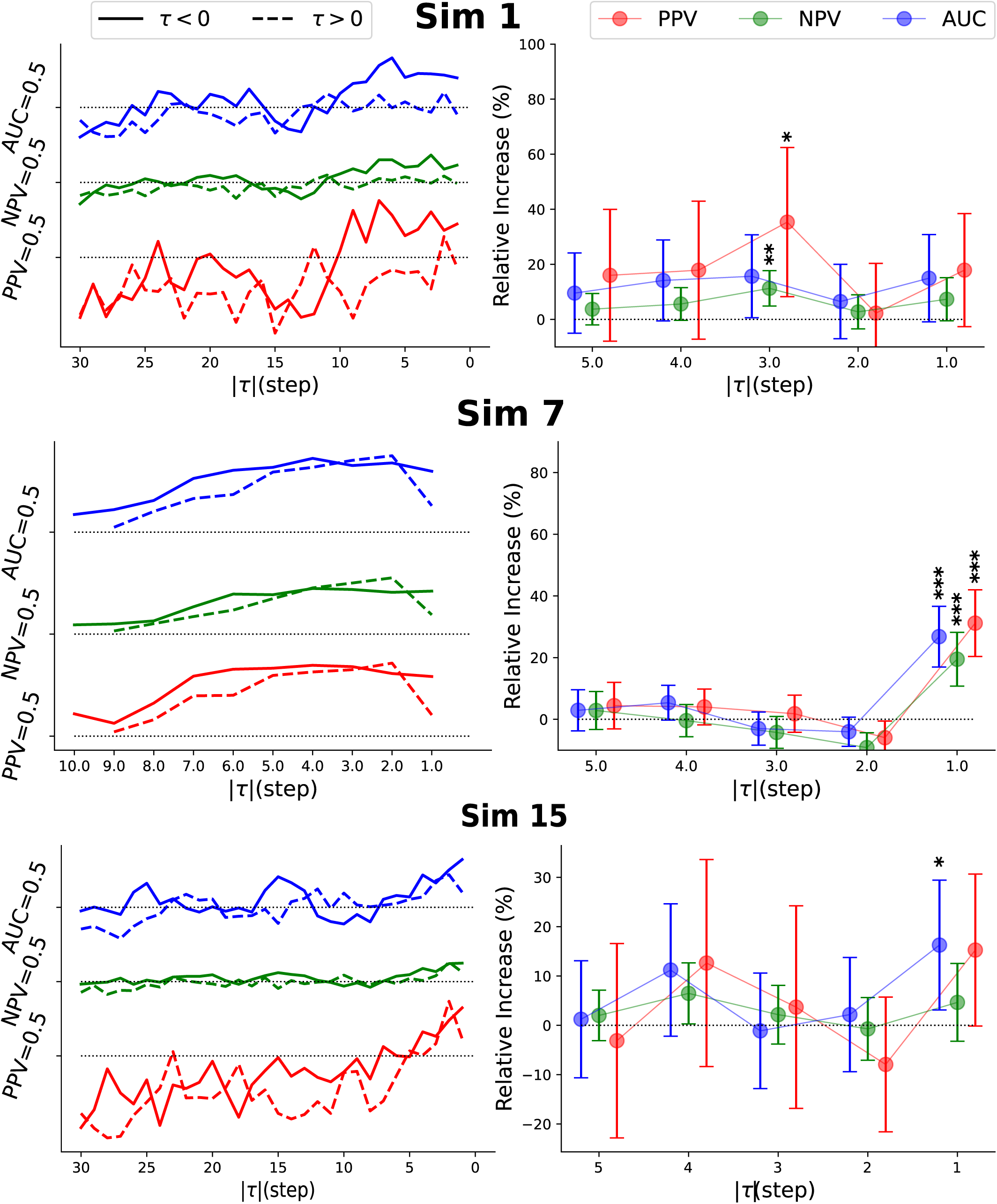
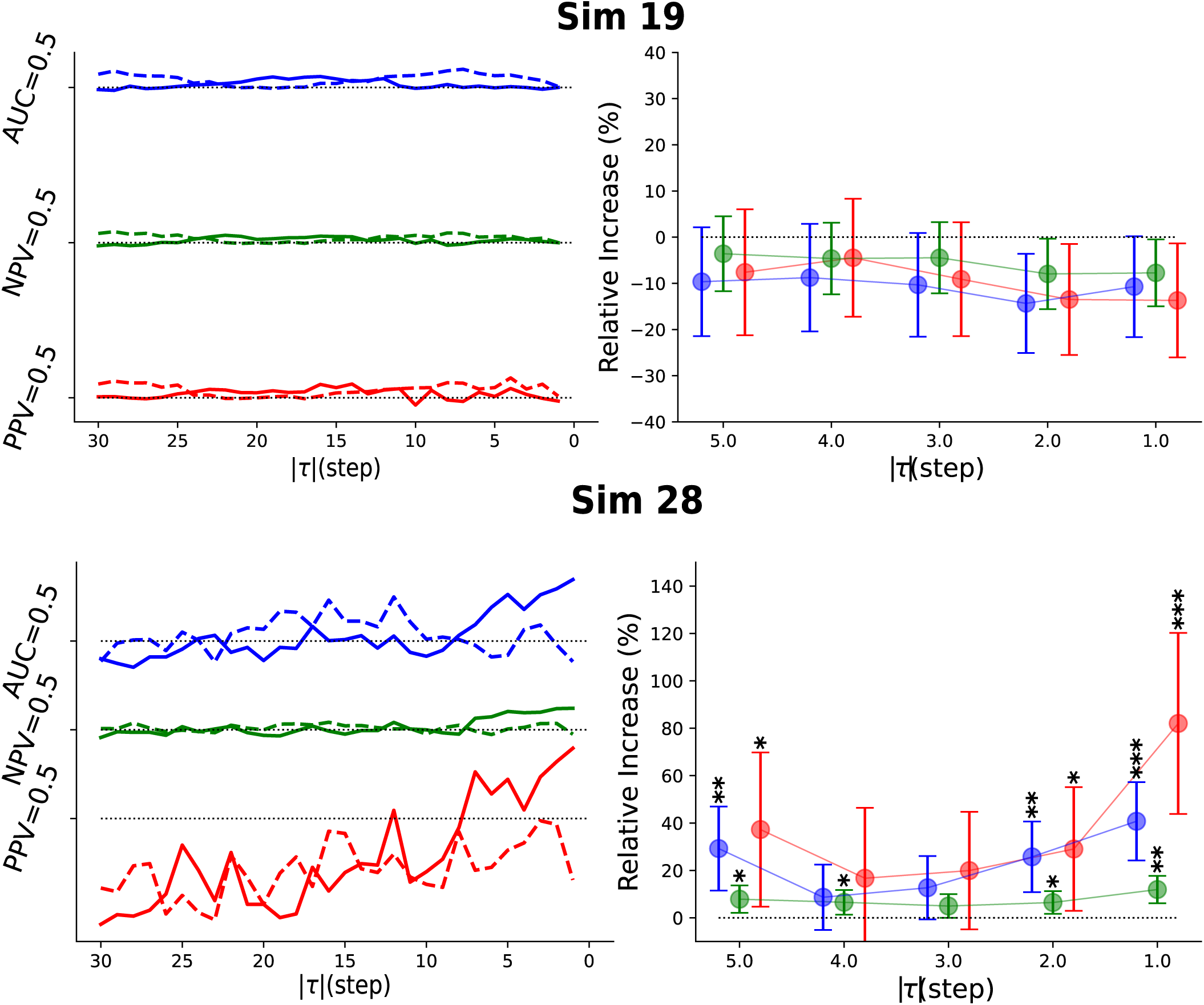
RCC metrics for negative and lags in the Netsim dataset. LEFT Mean metrics of RCC for each simulation tested for negative and lags. In all plots, the dashed black lines indicate the *better-than-chance* threshold. RIGHT Relative increase of metrics from RCC in the negative lag domain with respect to scores obtained with RCC in the positive lag domain for the first 5 lags accompanied with the corresponding significance analysis. Statistical significance was assessed using one-sided Welch’s t-tests given the hypothesized improvement of RCC in directionality detection (* for *p <* 0.05, ** for *p <* 0.01, and *** for *p <* 0.001).

### Consistent posterior-to-anterior interactions within the default mode network arising with RCC

We applied our framework to study whether coherent effective connectivity networks were recoverable. Given the slightly better predictive power of small and negative interaction delays (Fig. 6; see also Figs. S3 Figure to S7 Figure), we narrowed our analyses to a delay of 1 TR (i.e., *τ* = *−*2s). This is also consistent with most of the literature that considers large coupling delays to be highly unrealistic [18]. However, for transparency we report other times in the supplements. Subject-specific asymmetric adjacency matrices were obtained with Eq. 9. Each one of the two contributions was binarized based on its own significance threshold after correcting for the appropriate number of multiple hypotheses testing (see Materials and Methods). Importantly, we took a conservative approach and primed bidirectional interactions over directed ones to reduce the number of potential false positives - although the absence of any ground truth did not impose any clear restrictions. In the case were there was significant evidence for both bidirectional and one-directional interactions we disregarded the asymmetric component. Note how directionality scores are incompatible between opposite directions but the restriction on the bidirectional score is purely statistical. In practice this ment thresholding Eq. 9 and assigning to a particular connection either 0 or 1 based on the existence of at least one significant score. Finally, we studied the consistency of these binary directed networks across the entire population (*n*=223) by averaging the binary networks. In Fig. 7 we report connections with consistencies superior or equal to 0.75 (see S9 Figure for the actual numbers). We applied a gross division of posterior-temporal-anterior regions to simplify and compare the results with existing studies [10]. We found a high number of *intra*-modular symmetric connections and, noteworthy, a distinct pattern of asymmetry between anterior and parietal regions towards prefrontal cortex. Results were considerably robust across reasonable thresholds and the first TRs (see S10 Figure).

**Fig 7.**
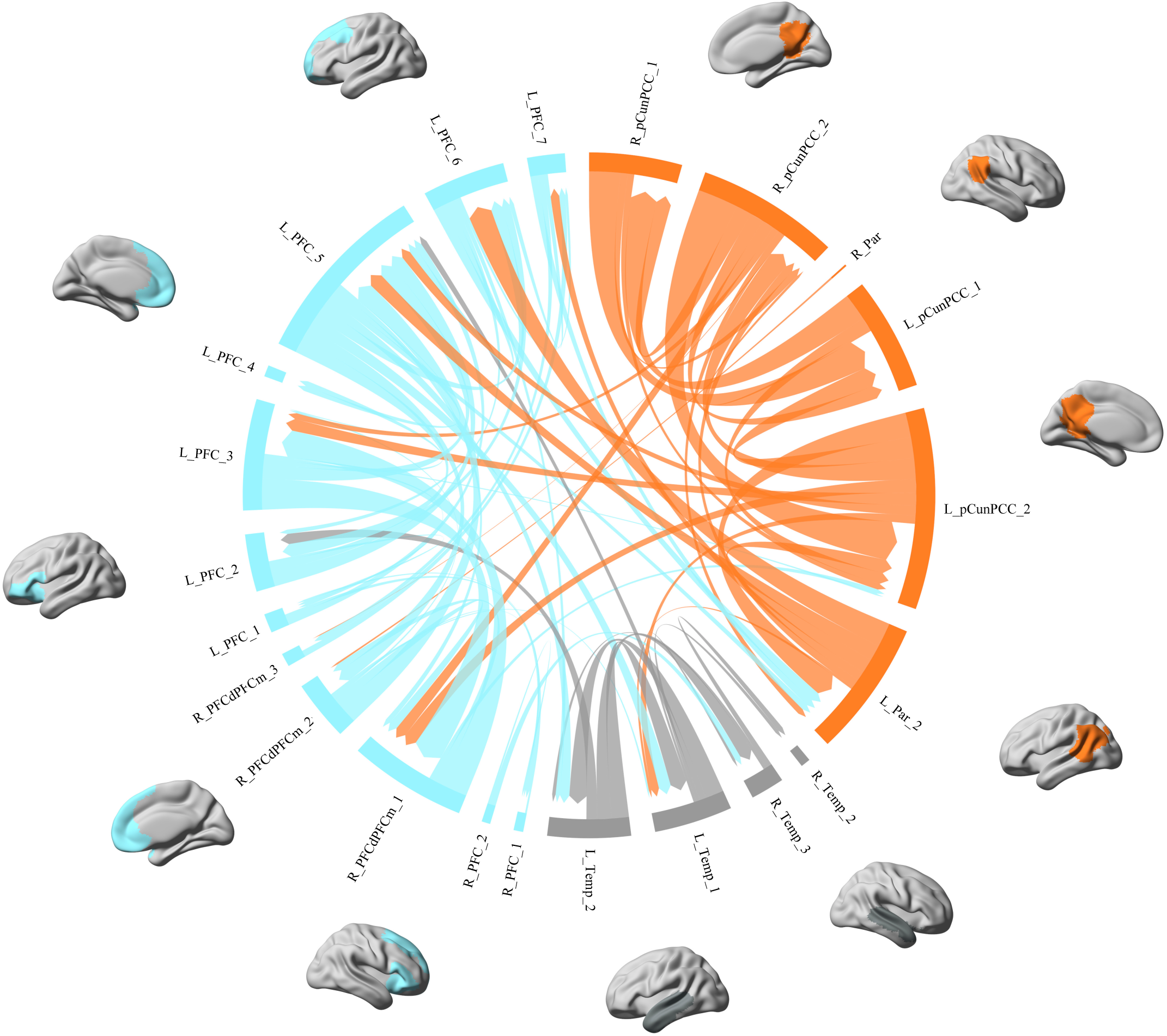
Most consistent effective connections within the default mode network. Directed connections with consistencies superior or equal to a 0.75 threshold across the entire population (*n*=223). Default mode network (DMN) regions were grouped into three categories based on anatomical and functional proximity. Light blue regions correspond to anterior regions - mainly prefrontal cortex (PFC), gray regions correspond to grossly the two hemispheric temporal lobes and, orange regions make for posterior regions including parietal and precuneus. A clear and distinct pattern of posterior-to-anterior influence is observable by means of orange edges terminating in blue regions but not the other way around. The thickness of the connections is proportional to the consistency of the connection across subjects.

## Discussion

Due to the intricate analysis of the results, the use of State Space Reconstruction (SSR) methods in neuroscience has so far been scarce [44] with inconclusive results [74]. Even more, in the specific case of Reservoir Computing Causality (RCC), we were not able to find earlier attempts to translate the method into network neuroscience. In a broader context, its use has been limited to small (sub-)systems [48]. The infancy of RCC might be the leading cause of this. We may argue that the main reason is the lack of a systematic method to assess the statistical validity of the directionalities inferred by the trained Reservoir Computing Networks (RCNs). Nonetheless, the lack of interpretability as opposed to its leading competitors, namely Granger Causality (GC) approaches, should also be regarded as one of its pitfalls. To this end, we presented a set of, hopefully, unambiguous steps that ultimately produced a reduced set of scores bounded between 0 and 1. These scores naturally incorporate statistical inference, therefore, making its interpretation formally equivalent to *p*-values and GC-based methods.

For the choice of reservoir parameters, we employed our method to disentangle directionality in a synthetic system that exhibited chaotic dynamics. Importantly, contrary to standard practice we did not simulate long series exempt of noise since we were interested in emulating realistic conditions. That is, we generated short and noisy sequences that would represent a challenge to any available method. Within these settings, the combination of RCC and the proposed analysis was able to successfully detect influence in a limited range of delayed effects. Notably, the predictive power was considered both in positive and negative lags.

We further tested our proposal in a set of fMRI simulations [73]. These simulations are known to be challenging to different causality detection methods thus being considered as reliable benchmarks. We evaluated the performance on a set of 5 simulations created to mimic different acquisitions. In contrast to other SSR methods the performance was independent of the length of time series, only showing sporadic linear trends. This was hinted at in the original proposal [48] but never properly explored. Moreover, the accuracy and the predictive powers of our proposal dropped to the *better-than-chance* threshold for long delays indicating that it was not learning spurious trends in the data. It is well-known that long delays are not biologically possible hence indicating that the trained RCNs were not learning nor introducing data-related artifacts.

When comparing these results with the outcome of GC, we performed systematically and, overall, significantly better in the negative lag domain. Given that GC can only test in the negative regime, we compared the performance of our proposal in the positive lag regime with the results in the negative domain. Contrary to what we expected, negative lags did not show clear improvement over positive delays. However, in some cases, the increase in classification scores was significantly higher on the negative side.

Based on these results, we studied resting-state fMRI data within the time domains where results from the synthetic data were reliable and consistent with the literature [18, 73]. Within the DMN, there is a distinctive pattern of symmetry that characterizes its functional connectivity. Indeed, studies using fMRI have revealed that the DMN exhibits a predominantly bidirectional flow of information between its constituent regions. However, even in the absence of a ground-truth, there have been significant attempts to address causality within the DMN. Initially, it was suggested that the core DNM regions exhibit stronger influence from anterior (e.g., medial prefrontal cortex) to posterior regions (e.g., precuneus and posterior cingulate gyrus) [9]. However, more recent studies tend to indicate the opposite pattern among medial regions [11, 65, 68]. This pattern was also demonstrated between lateral posterior and anterior regions [10, 66, 68]. Yet, the discrepancy is still notable [66, 67], probably due to the lack of task demands during resting-state conditions. In accordance with these observations, our results indicate some bidirectional interactions, however, with a more prominent influence from posterior to anterior regions. Higher consistency appeared in connections that were originating in posterior areas and terminating in anterior regions (Fig. 7).

Furthermore, the driving role of left parietal regions was found to be more prominent than its contra-lateral homologous as found in several studies [10, 11]. Temporal lobes seemed to be less connected with the rest of the resting-state network, although some connections emerged from the temporal and terminated in frontal regions. This is consistent with the role of medial anterior regions (e.g., mPFC) in integrating sensory information [75], although it is unclear what information would be integrated during resting conditions.

### Alternative proposals based on RCNs

There are several advantages of using RCNs over other machine learning frameworks for quantifying effective connectivity across macroscale functional interactions in the brain. Reservoir-based systems can handle stochastic, deterministic, linear, and non-linear dynamics [76] while bridging the gap between dynamical and statistical approaches [77]. The most straightforward way to leverage the forecasting power of RCNs is to substitute the autoregressive model with an RCN. It has been shown that the results can be interpreted identically as in canonical GC [78]. Alternatively, an analogous statistical score can be defined to determine directionality [79]. Despite inheriting the known conceptual drawbacks from GC-based approaches [18], interestingly, this simple substitution, yielded reduced false positive rates in certain *in silico* datasets. However, it can be argued that the improvement over RCC is rather limited since the RCN architectures used in [79] significantly differed from the original proposal in [48] and in this work. Quoting the authors: ”high false positive for RCC is mainly due to one-reservoir in RCC”. Hence, RCC might be, at least, equivalent if more complex architectures were studied.

Although not presented here, we explored more intricate RCN architectures for the *noise-free* logistic maps in Eqs. (11) and Fig. 2 without observing improved results. This may originate in the initial optimal performance in identifying directionality based on single reservoirs. When considering the rest of the use cases explored in this work, limited data points, large sample sizes, and the computational demands of the process we developed made the exploration of these more complex architectures problematic. Nonetheless, we did implement the possibility of trying out both sequential and parallel reservoir blocks that could potentially increase the predictive capacity of the RCNs and the whole RCC rationale (see Data and code availability). Ultimately, one would hope that, given the adequate data needed for these kinds of studies, the flexibility of PyRCN [50] would allow for more rigorous testing and exploration of the proposed method and/or variants. Consequently, the development of a systematic approach for evaluating causal interactions within an RCC framework represents a necessary step toward achieving a reliable RCC detection ecosystem.

A similar idea was recently explored by Suzuki and colleagues [77]. Instead of optimizing predictabilities by fine-tuning a single reservoir architecture, they tackled the problem of which series are *valuable* predictors of the desired target. In this regard, their method falls again into a Granger-based paradigm where the measure of directed influence relies on the extent to which a consequence is predicted by a discrete set of causes. For this purpose, a fast adaptive method was developed that automatically incorporates and/or removes potential predictors of the selected target as input to a set of RCNs by iterating through all the possible edges in the network. Of particular relevance is the fact that the number of additions and removals scaled with the number of nodes, making its direct applicability to large brain networks challenging. However, similar constraints or priors as the ones we use might prove useful when using this proposal. Their application was on ecosystem networks, which, despite their considerable size were not comparable to standard brain parcellations [64]. An interesting claim not present in [79] is that this iterative strategy can disentangle chained (e.g., *X → Y → Z* as well as fan-out events (e.g., *Z ← X → Y*), yet, RCC has been proven to reconstruct similar relationships [79] (see also Fig. 2).

### On the nature of Reservoir Computing Causality

We would now like to emphasize the paradigm shift that RCC might represent against Granger-based modeling. According to Granger [80], future events are caused by previous ones, simultaneous, or both; but the inverse relationships are not possible. This assumption was recently challenged [81] by showing that a reversed time series is an equally valid predictor of the future. Contrary to this particularly concerning flaw, CCM-based reasoning, of which RCC is part of, is not insensitive to the temporal structure of the system (Fig. 8). Although we lack formal proof of the reason why this is true, we are inclined to believe that this is due to the significant change of the attractor’s landscape that the reversal of one of the series causes. Therefore, under these circumstances, the applicability of Taken’s theorem, necessary for the existence of CCM-causality, is not guaranteed (see S8 Figure).

**Fig 8.**
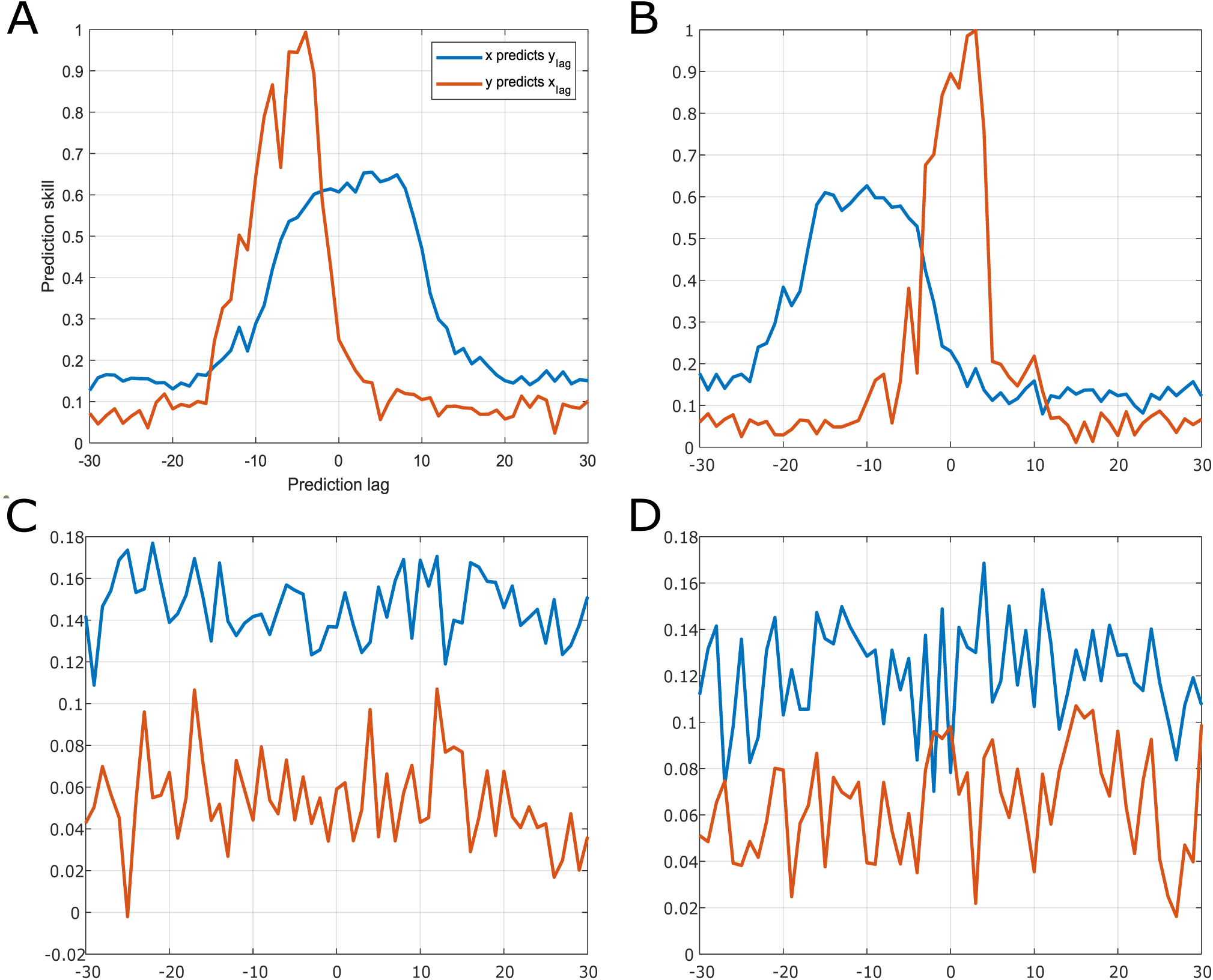
Temporal asymmetry of RCC. Predictability scores from the system defined in Eq. 11 and [48]. The codes from the original publication were used with consent from the authors. **A** The original time series. **B** Both time series were temporally reversed. **C** Only time series *x_t_* was reversed, that is, *x_t_ → x_−t_*. **D** Only the time series *y_t_* was reversed using the same procedure. Note how only in the case where both time series are reversed can an optimal mapping between shadow manifolds be found.

The scores we proposed in this work consider the evidence concerning one direction together with the non-existence of evidence in favor of the opposite direction. Furthermore, we independently considered symmetrical relationships from the individual interactions between the two directions. That is, we defined an independent score considering the lack of evidence for any directed link hence only monitoring mutual influences. Note, however, how in GC-based approaches bidirectionality is assessed separately by the evidence for each direction. In summary, as opposed to GC-based methods, the analysis we proposed considers the convergence of multiple sources of statistical knowledge to define whether 1) an asymmetric influence can be considered independently, 2) the data supports symmetric interactions, or 3) the absence of any sort of coupling.

### Limitations and future work

Further and improved inferences from RCC might come from the use of newer machine learning systems. In this regard, new generation RCNs [82], deeper combinations of deep and reservoir learning [50] and increased reliability in fMRI acquisition [83] show great potential to unravel a detailed effective map of the human brain [84].

Another interesting yet non-trivial research line in neuroscience aims to understand behavior through state space analysis and neural manifolds [37, 85]. Given that fMRI time series contain behavioral correlates, it is not unlikely that analysis of the internal states of the trained RCNs might shed some light on the sequence of events hidden in the data. It has been recently shown how transitions from brain states are mediated by the attractor space, which, hopefully, would be captured by the reservoirs.

Finally, the chances of spurious causalities and unseen variables can never be completely ruled out. Inferences from model-based (e.g., causal modeling) are often interpreted as closer to the natural relationships one wishes to reveal. Yet, issues such as the prohibitive size of the possible models or the computational constraints of the chosen model remain. On the other hand, model-free approaches offer an appealing solution but, as we have tried to emphasize throughout this manuscript, their results should be interpreted carefully. To follow more carefully causal brain dynamics, modulation by neurostimulation has been proposed [86]. Specifically, by perturbing a source region through neuromodulation and observing the response in the brain. This is challenging for many reasons from the need to have a neurostimulation device, to concurrent stimulation and monitoring. Similarly as done in a work using multivariate Ornstein-Uhlenbeck processes [19], computational approaches can be used to define a surrogate brain and then implement systematic perturbations as selective increases of the BOLD signal at specific regions. By observing the induced responses, perturbation-based effective connectivity can be inferred. However, we argue that the construct of the surrogate brain does not necessarily invalidate the issue of spurious temporal correlations. Instead, efforts need to be taken to define and fit the surrogate model to avoid reiterating the same conceptual drawbacks [84]. This further validation and incorporation into RCC is beyond the scope of the current work and remains to be explored.

## Conclusion

We have presented a systematic approach to understanding and analyzing the outcomes of reservoir computing networks trained to discriminate directed relationships in a set of challenging dynamics systems. We provided a comprehensive use case of reservoir computing causality and showed its improved inferences compared to Granger causality. We tested our proposal in fMRI simulations reaching better accuracies and reconstructed a directed resting-state network derived from scanned human brain activity. Overall, this work represents a promising first step toward the establishment of alternative but reliable brain causality methods.

## Acknowledgments

This research is supported by the European Union’s Horizon 2020 research and innovation programme under grant agreement Sano no 857533, and by the International Research Agendas programme of the Foundation for Polish Science, co-financed by the European Union under the European Regional Development Fund. This research was supported in part by the PLGrid infrastructure. Computations have been performed at Ares supercomputer at ACC Cyfronet AGH. We also want to thank Alicja Olszewska and Filip Sondej for the useful insight during the Krakow Brainhack 2022.

## Author contributions

**Conceptualization** Joan Falćo-Roget, Adrian I. Onicas, Felix Akwasi-Sarpong, Alessandro Crimi

**Data Curation** Joan Falćo-Roget, Adrian I. Onicas

**Formal analysis** Joan Falćo-Roget, Adrian I. Onicas

**Investigation** Joan Falćo-Roget, Adrian I. Onicas

**Methodology** Joan Falćo-Roget, Adrian I. Onicas, Alessandro Crimi

**Project administration** Alessandro Crimi

**Software** Joan Falćo-Roget

**Supervision** Alessandro Crimi

**Visualization** Joan Falćo-Roget, Adrian I. Onicas

**Writing - Original draft** Felix Akwasi-Sarpong, Alessandro Crimi

**Writing - review & editing** Joan Falćo-Roget, Adrian I. Onicas, Alessandro Crimi

## Supplementary Material

## S1 Appendix Heuristics to define the directionality scores

- Consider an example of **x** *→* **y** tested in the ***positive*** *τ* regime. The minimum things required to consider directionality are the following:

1. a statistically significant predictability of the **y** series from **x** with respect to its surrogate relationship and,
2. a statistically significant evidence for positive Δ-scores with respect to the surrogate score; specifically, the Δ-score computed using the predictabilities of both surrogates **x***_s_* and **y***_s_*.
- Consider an example of **y** *→* **x** tested in the ***negative*** *τ* regime. The minimum things required to consider directionality are the following:

1. a statistically significant predictability of the **y** series from **x** with respect to its surrogate relationship and,
2. a statistically significant evidence for positive Δ-scores with respect to the surrogate score; specifically, the Δ-score computed using the predictabilities of both surrogates **x***_s_* and **y***_s_*.
- Consider an example of **y** *→* **x** tested in the ***positive*** *τ* regime. The minimum things required to consider directionality are the following:

1. a statistically significant predictability of the **x** series from **y** with respect to its surrogate relationship and,
2. a statistically significant evidence for negative Δ-scores with respect to the surrogate score; specifically, the Δ-score computed using the predictabilities of both surrogates **x***_s_* and **y***_s_*.
- Consider an example of **x** ⇄ **y** tested in both the ***negative*** and ***positive*** *τ* regimes. The minimum things required to consider directionality are the following:

1. a statistically significant predictability of the **x** series from **y** with respect to its surrogate relationship and,
2. a statistically significant predictability of the **y** series from **x** with respect to its surrogate relationship and,
3. a ***non***-statistically significant evidence for negative and positive Δ-scores with respect to the surrogate score; specifically, the Δ-score computed using the predictabilities of both surrogates **x***_s_* and **y***_s_*.

Note how the latter score evolves contrary to all the previous scores due to the non-significance of the Δ-score; that is, if they are different from zero (no matter which direction) evidence for bidirectionality would decrease due to small expected value of *p*_Δ_*_τ_*_(**x,y**)_*_̸_*_≠0_.

**S1 Fig .**
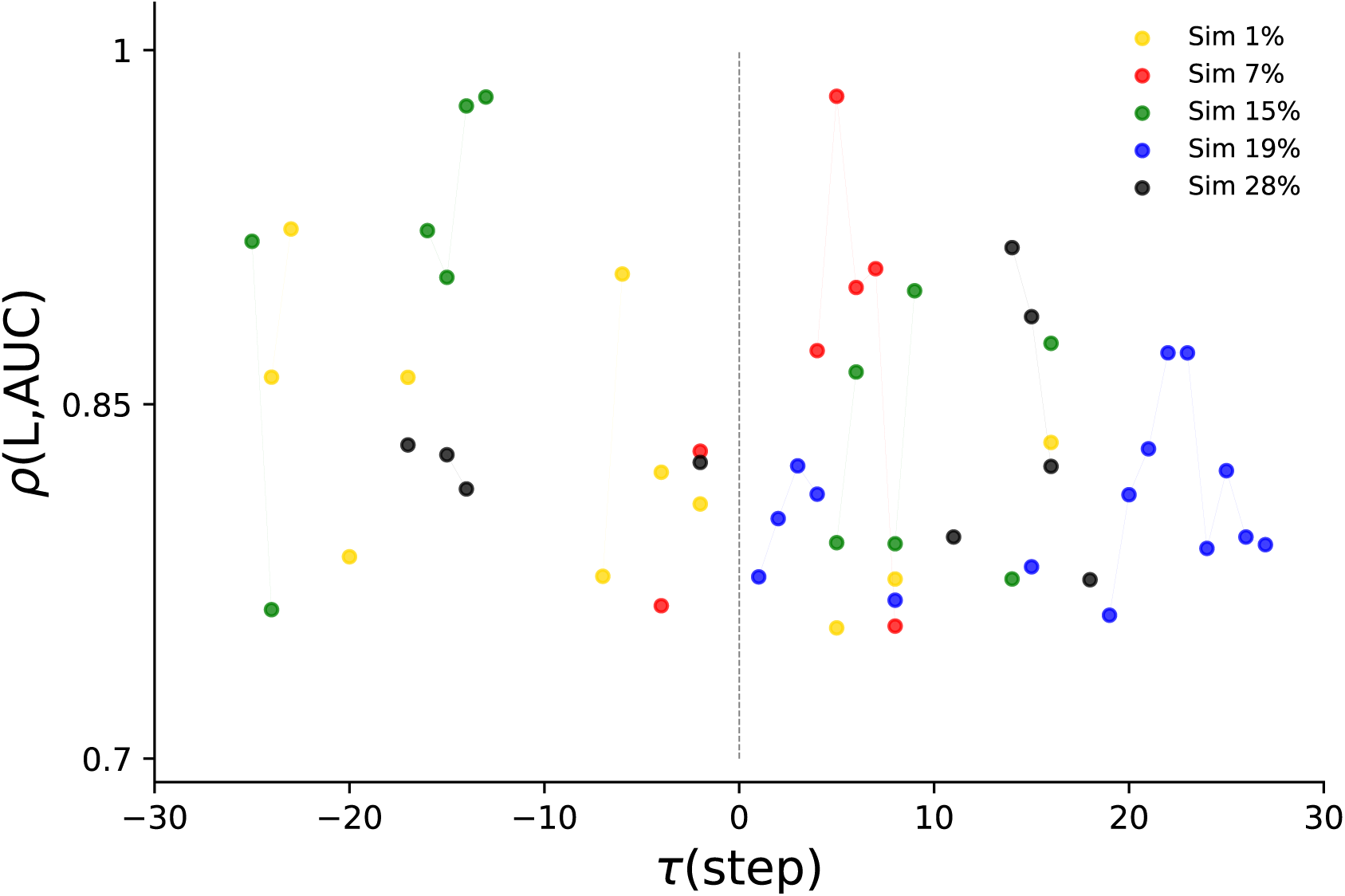
Correlations between time series length and classification accuracy in RCC. For each tested lag *τ*, we computed the correlation coefficient between the accuracy and the time series length (as described in the main text). We show filtered values (i.e., *ρ >* 0) that were statistically significant (*p <* 0.05, exact test). No clear nor consistent dependence between the accuracy of RCC and the length of the time series was found.

**S2 Fig .**
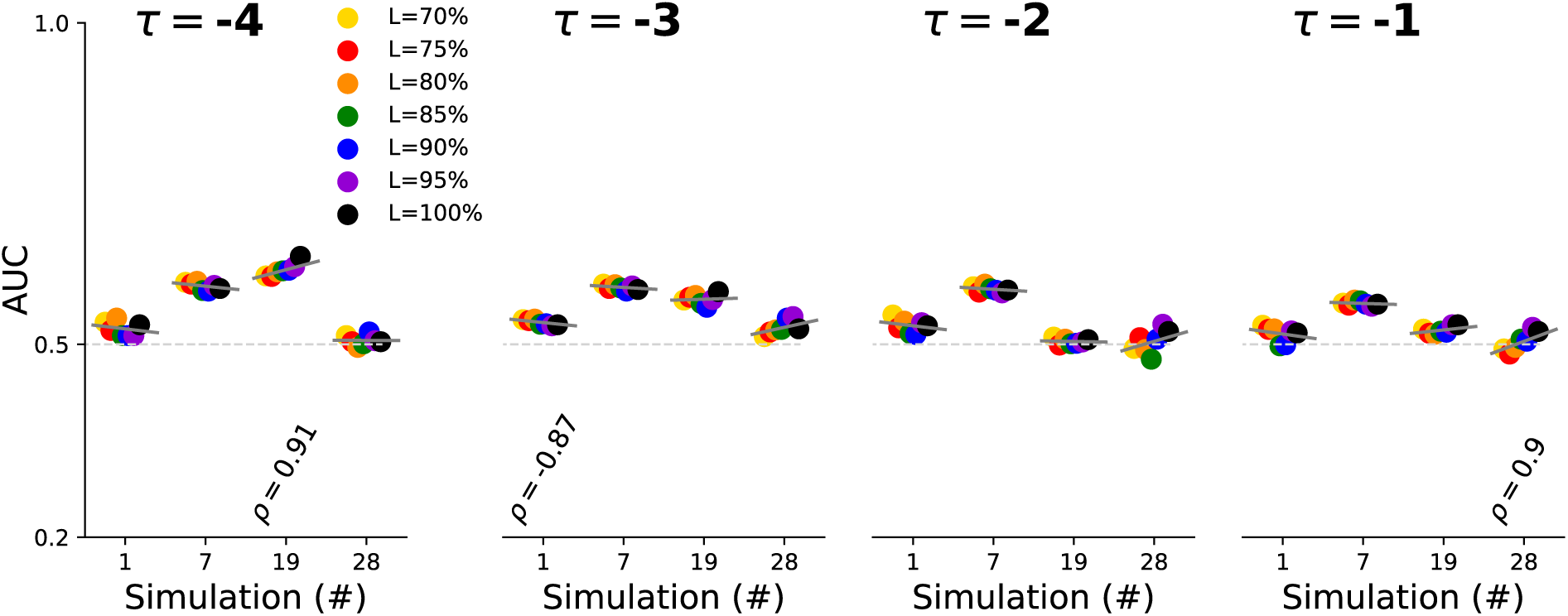
GC’s Convergence of the classification metrics as a function of the time series length. Area Under the receiver operating Characteristic (AUC) for each simulation and the first 4 selected lags *τ* for each one of the simulations tested in this work. Short gray lines depict linear fits between AUCs and time series length. Numbers show the values of the statistically significant correlation coefficients found (*p <* 0.05, exact test). Error bars are not shown for the sole purpose of clarity.

**S3 Fig .**
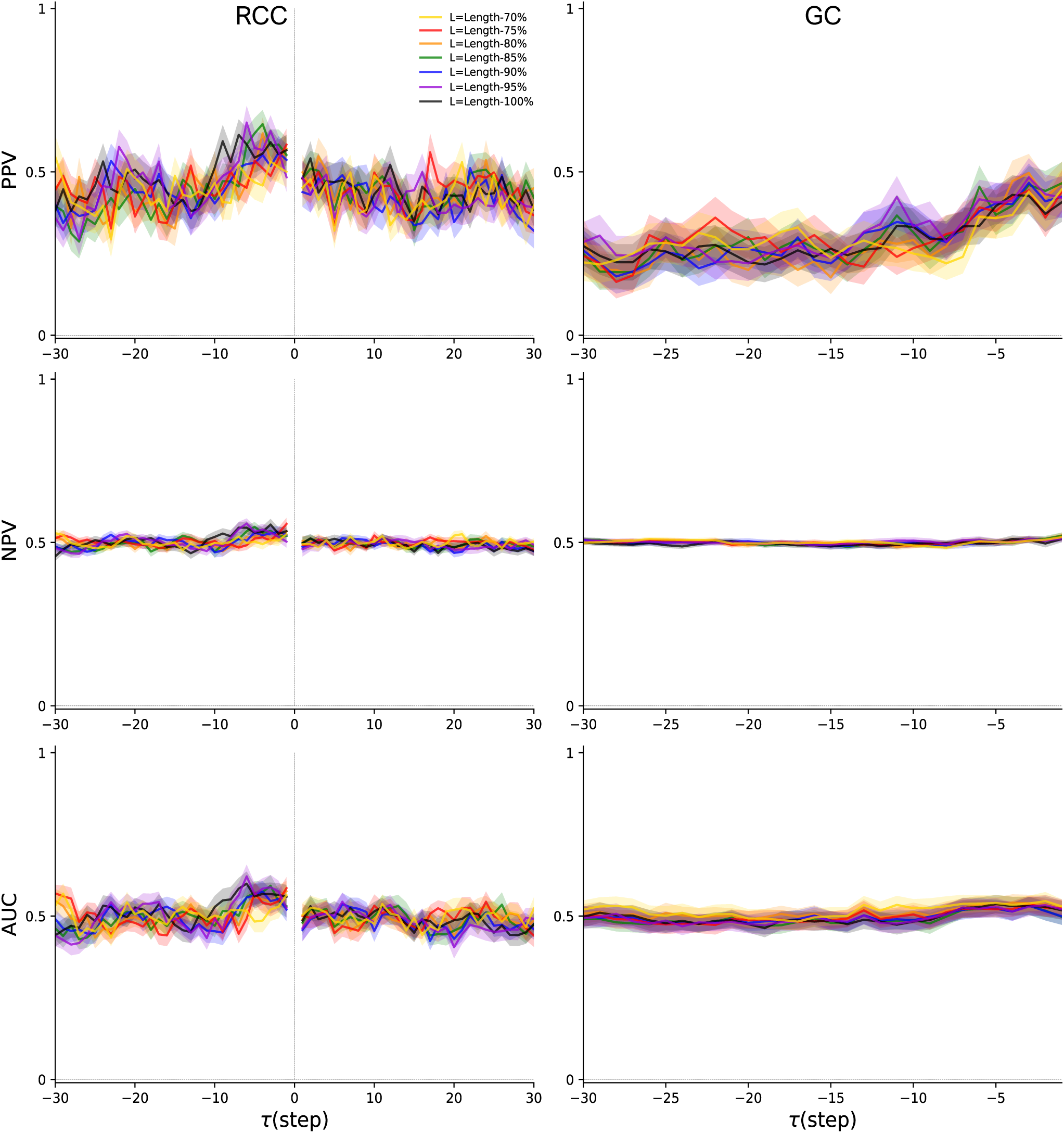
Directionality detection metrics for Simulation number 1 in the Netsim dataset. Results derived from Reservoir Computing Causality (RCC) in the left column and from Granger Causality (GC) in the right column. The metrics shown are Positive Predictive Value (PPV), Negative Predictive Value (NPV), and Area Under the receiving operating Characteristic (AUC). For both RCC and GS, the scores were obtained following the procedure described in the main text. Note how GC is only tested in the negative lag regime due to the original definition of it, hence making it not directly comparable to RCC results. All solid and shaded curves indicate results from the population analysis [mean*±*SEM].

**S4 Fig .**
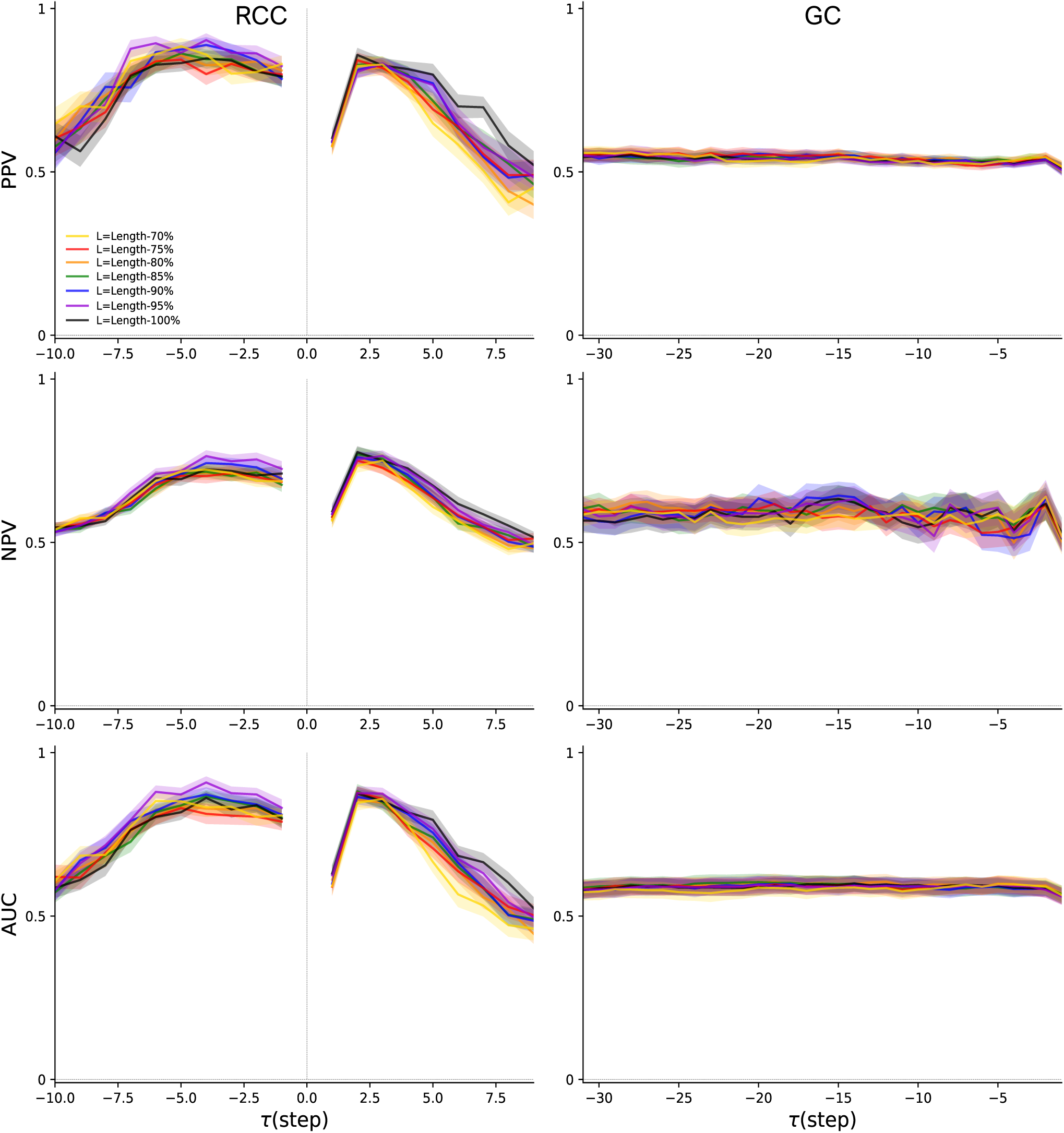
Directionality detection metrics for Simulation number 7 in the Netsim dataset. Results derived from Reservoir Computing Causality (RCC) in the left column and from Granger Causality (GC) in the right column. The metrics shown are Positive Predictive Value (PPV), Negative Predictive Value (NPV), and Area Under the receiving operating Characteristic (AUC). For both RCC and GS, the scores were obtained following the procedure described in the main text. Note how GC is only tested in the negative lag regime due to the original definition of it, hence making it not directly comparable to RCC results. The shorter range of tested lags for RCC was due to increased computation time. All solid and shaded curves indicate results from the population analysis [mean*±*SEM].

**S5 Fig .**
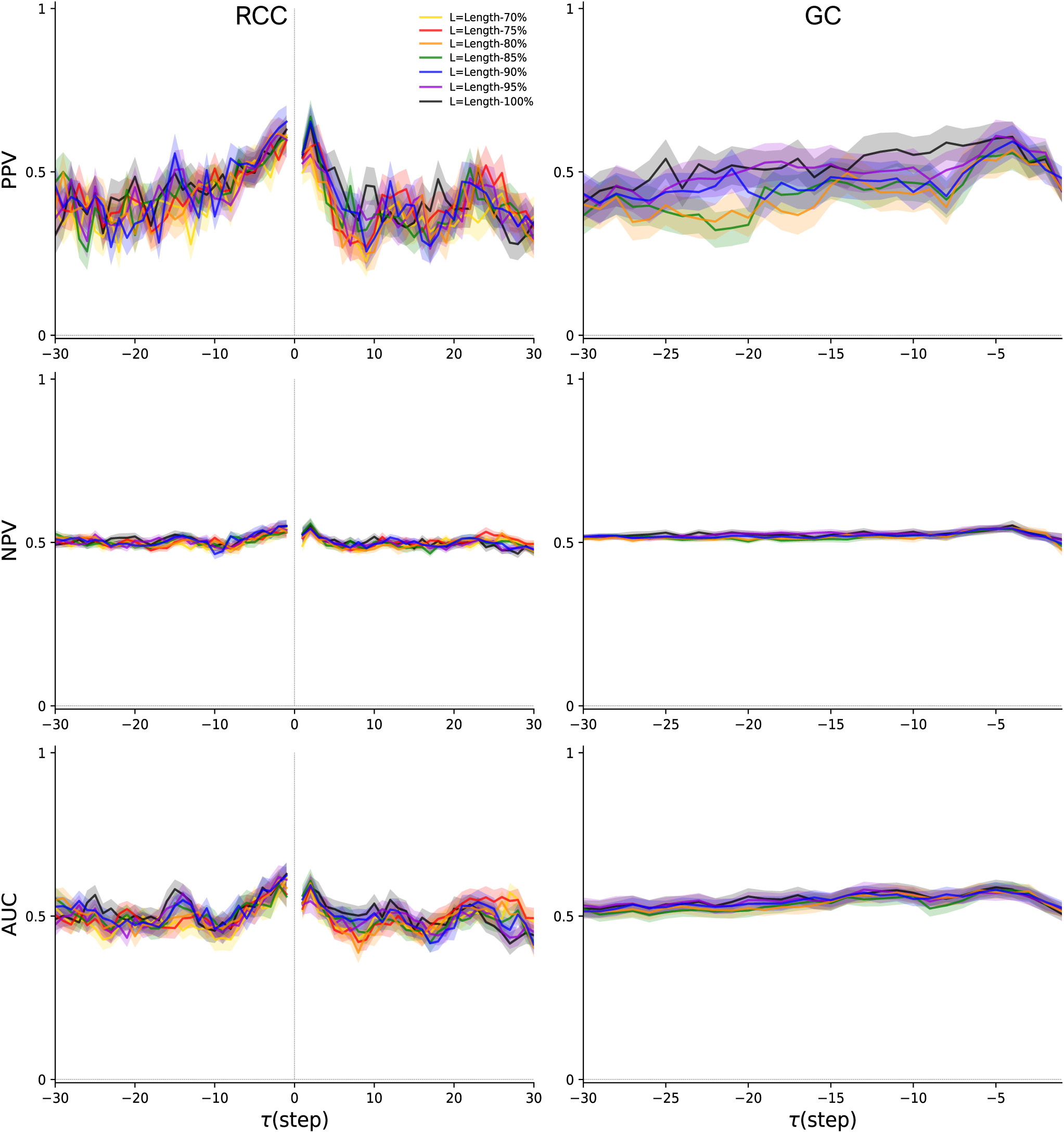
Directionality detection metrics for Simulation number 15 in the Netsim dataset. Results derived from Reservoir Computing Causality (RCC) in the left column and from Granger Causality (GC) in the right column. The metrics shown are Positive Predictive Value (PPV), Negative Predictive Value (NPV), and Area Under the receiving operating Characteristic (AUC). For both RCC and GS, the scores were obtained following the procedure described in the main text. Note how GC is only tested in the negative lag regime due to the original definition of it, hence making it not directly comparable to RCC results. All solid and shaded curves indicate results from the population analysis [mean*±*SEM].

**S6 Fig.**
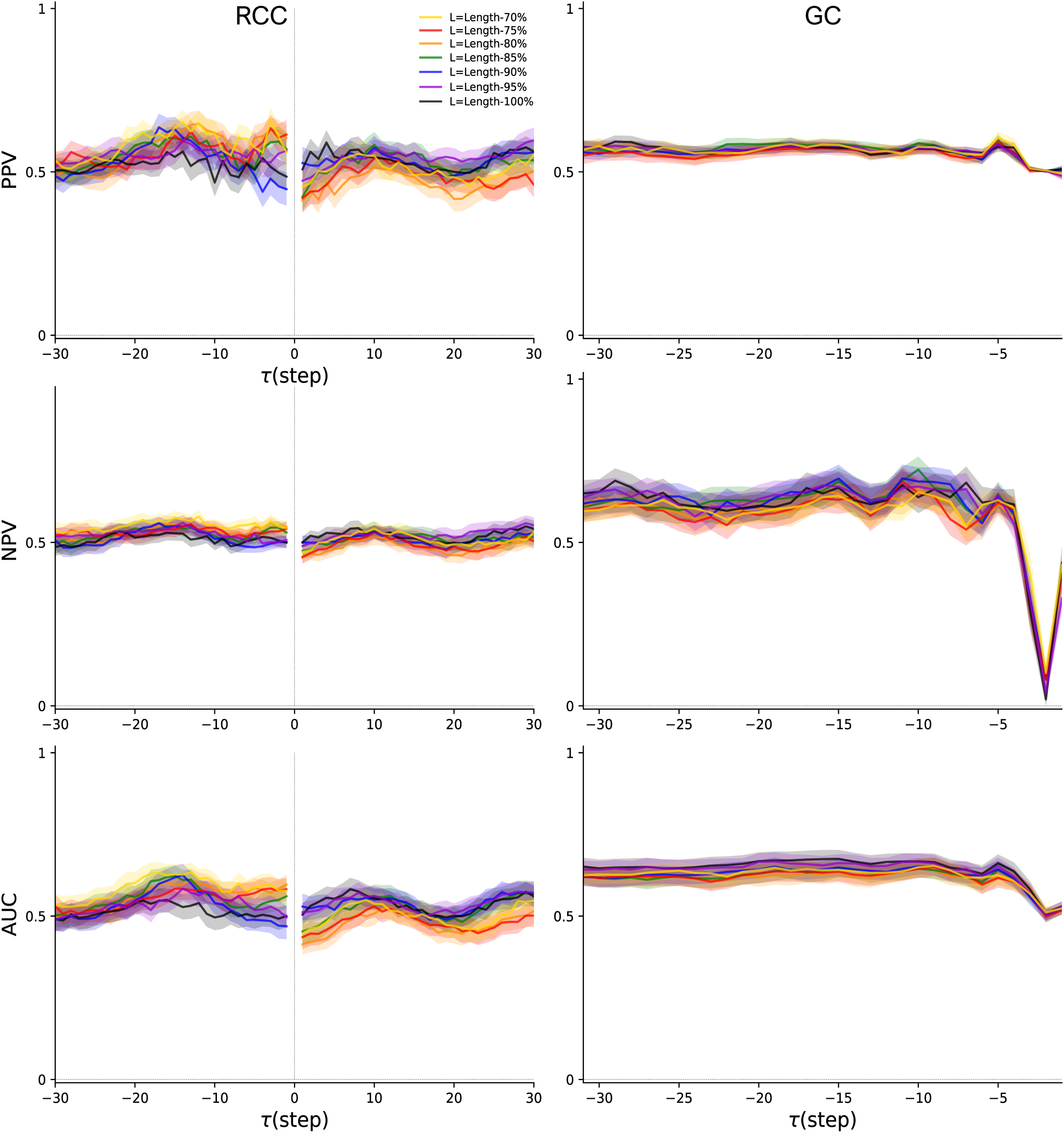
Directionality detection metrics for Simulation number 19 in the Netsim dataset. Results derived from Reservoir Computing Causality (RCC) in the left column and from Granger Causality (GC) in the right column. The metrics shown are Positive Predictive Value (PPV), Negative Predictive Value (NPV), and Area Under the receiving operating Characteristic (AUC). For both RCC and GS, the scores were obtained following the procedure described in the main text. Note how GC is only tested in the negative lag regime due to the original definition of it, hence making it not directly comparable to RCC results. All solid and shaded curves indicate results from the population analysis [mean*±*SEM].

**S7 Fig.**
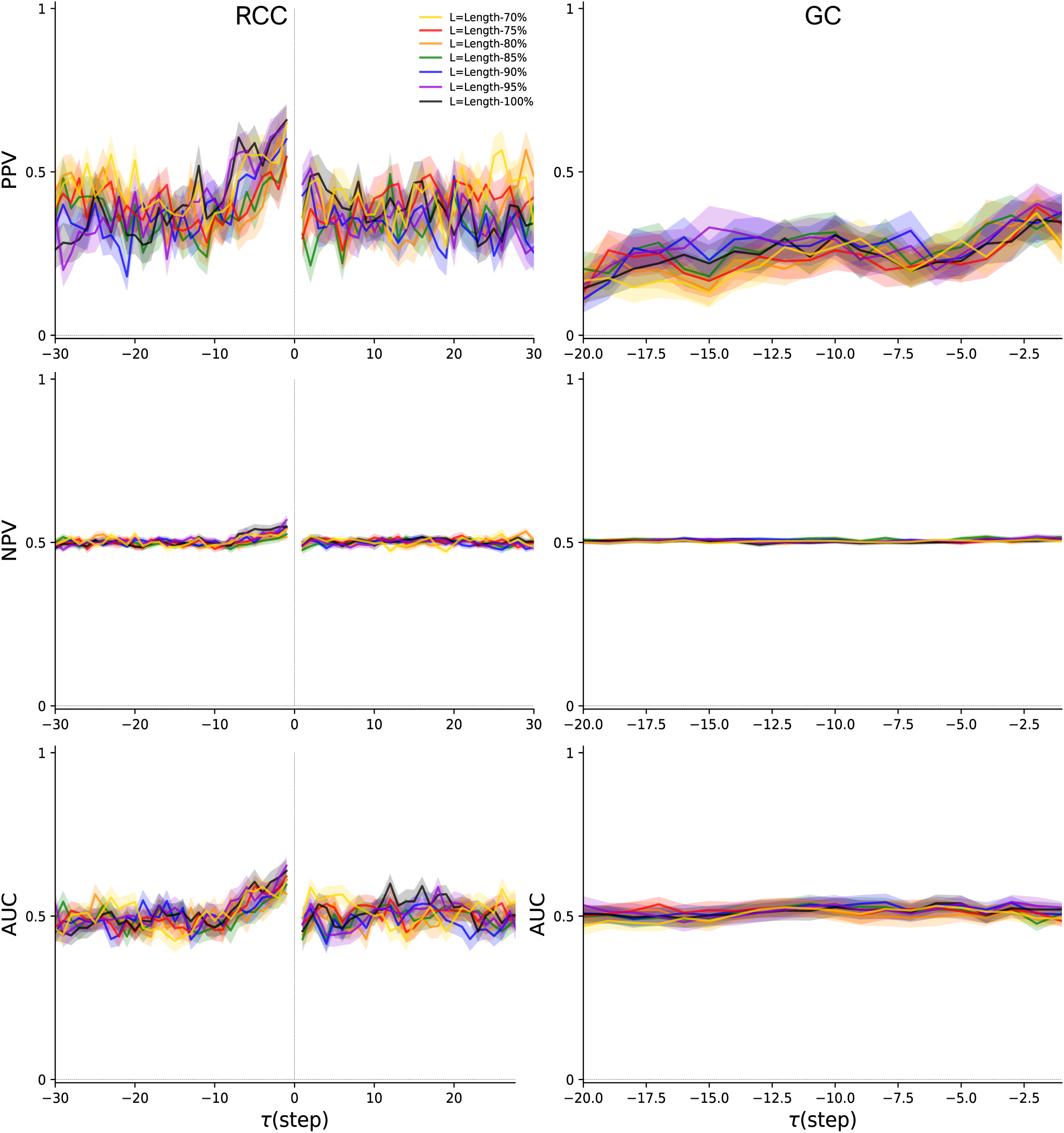
Directionality detection metrics for Simulation number 28 in the Netsim dataset. Results derived from Reservoir Computing Causality (RCC) in the left column and from Granger Causality (GC) in the right column. The metrics shown are Positive Predictive Value (PPV), Negative Predictive Value (NPV), and Area Under the receiving operating Characteristic (AUC). For both RCC and GS, the scores were obtained following the procedure described in the main text. Note how GC is only tested in the negative lag regime due to the original definition of it, hence making it not directly comparable to RCC results. All solid and shaded curves indicate results from the population analysis [mean*±*SEM].

**S8 Fig .**
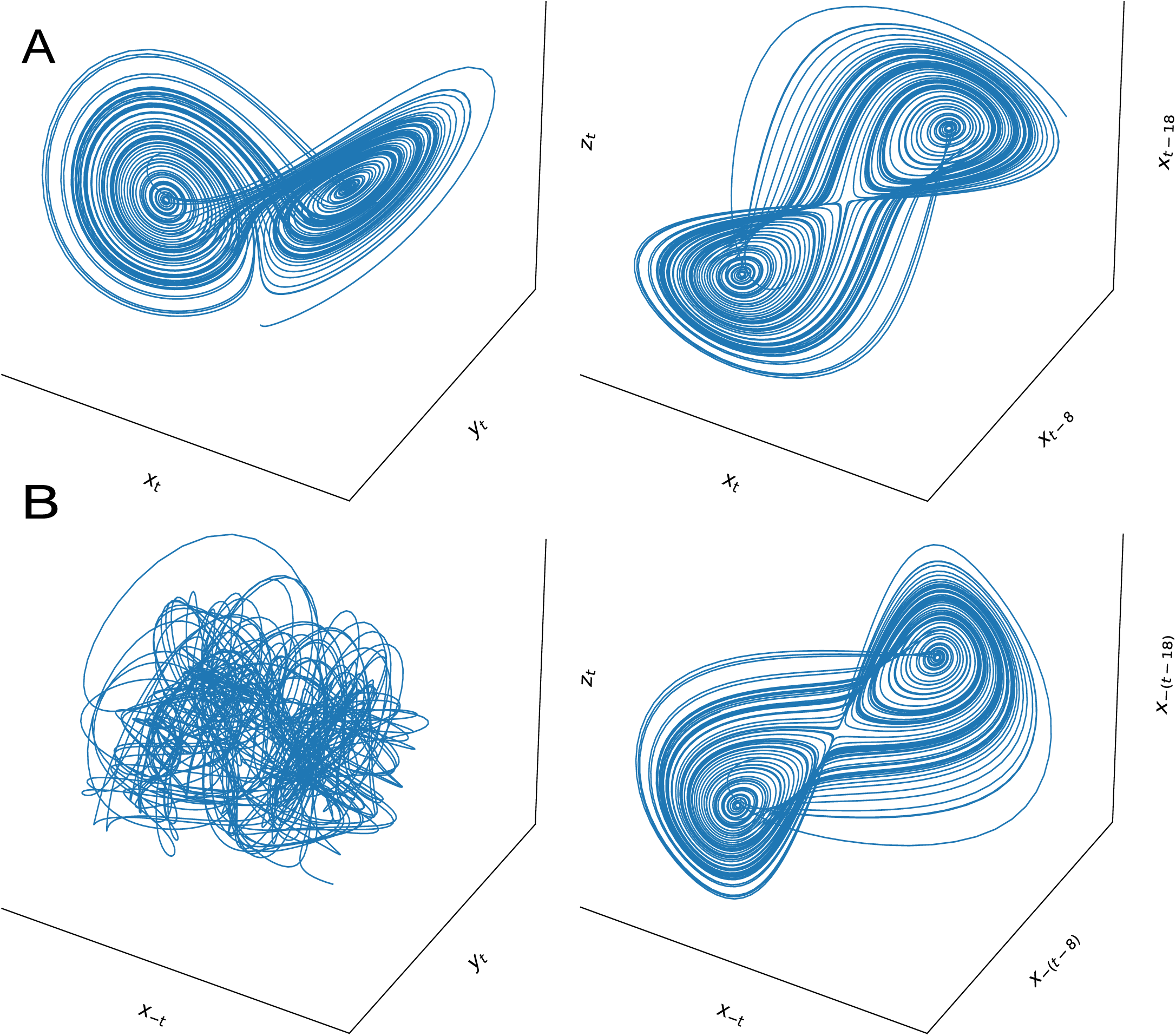
Lorentz attractor in reversed series. **A** State space of the original Lorentz system in the LEFT and the corresponding shadow manifold in the RIGHT. **B** State space of the Lorentz system after reversing one of the series (i.e, *x_t_ → x_−t_*) in the LEFT and the corresponding shadow manifold of the same variable in the RIGHT. Note how the respective manifolds built from the delayed series remain unaltered (except for a symmetric flip) but rather the original attractor has changed its topology and, in this case, completely vanished.

**S9 Fig .**
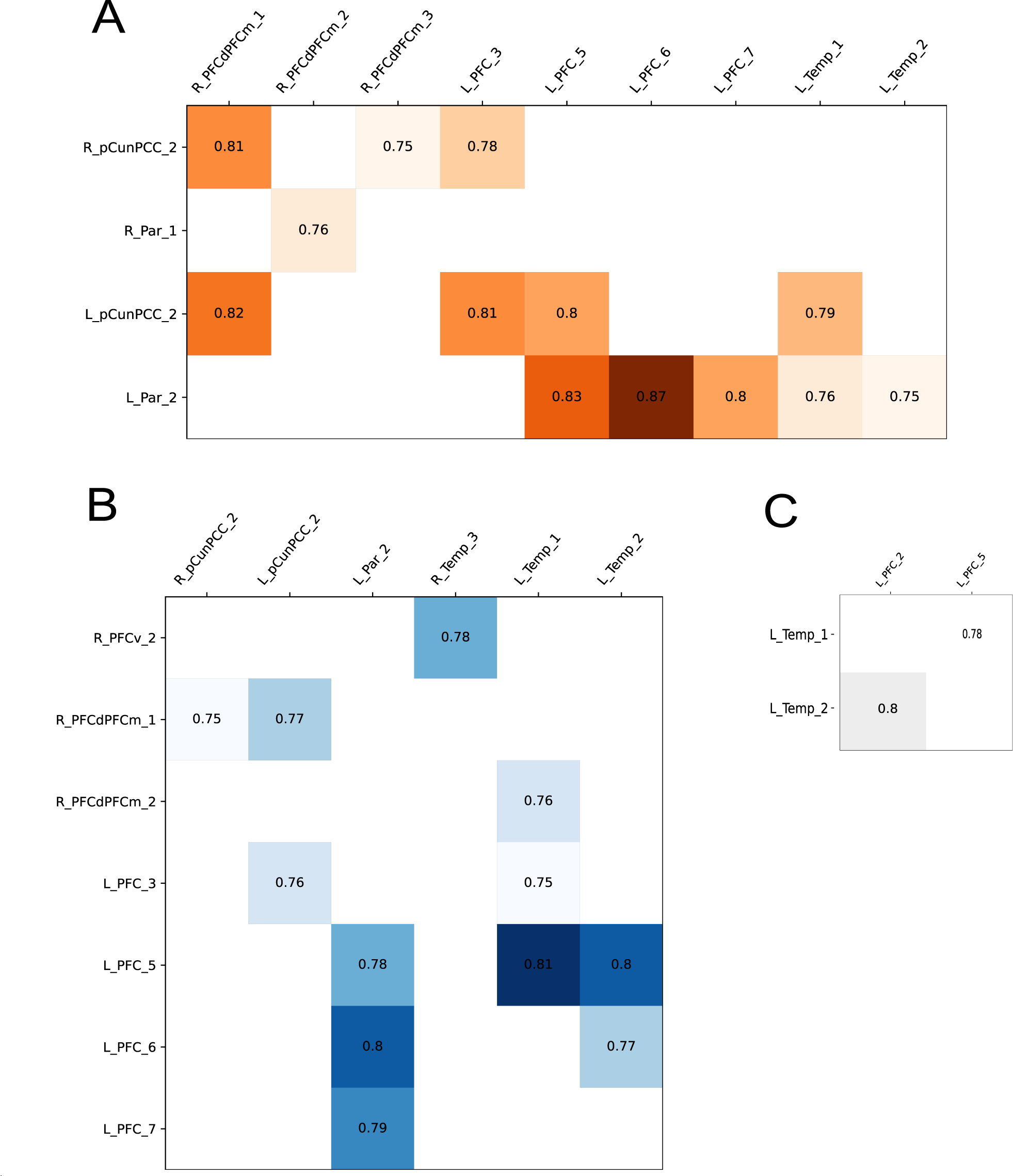
Numerical results for consistency across subjects. Consistency values above 0.75 for all the connections originating in each one of the three groups described in the main text. **A** Connections originating in posterior regions (rows) and terminating in anterior or temporal regions (columns). **B** Connections originating in anterior regions and terminating in posterior or temporal regions. **C** Connections originating in temporal regions and terminating in anterior or posterior regions. In this latter case, the number of consistent connections originating in temporal lobes was small and only terminated in two prefrontal regions. White connections did not exceed the 0.75 consistency threshold and are left empty.

**S10 Fig .**
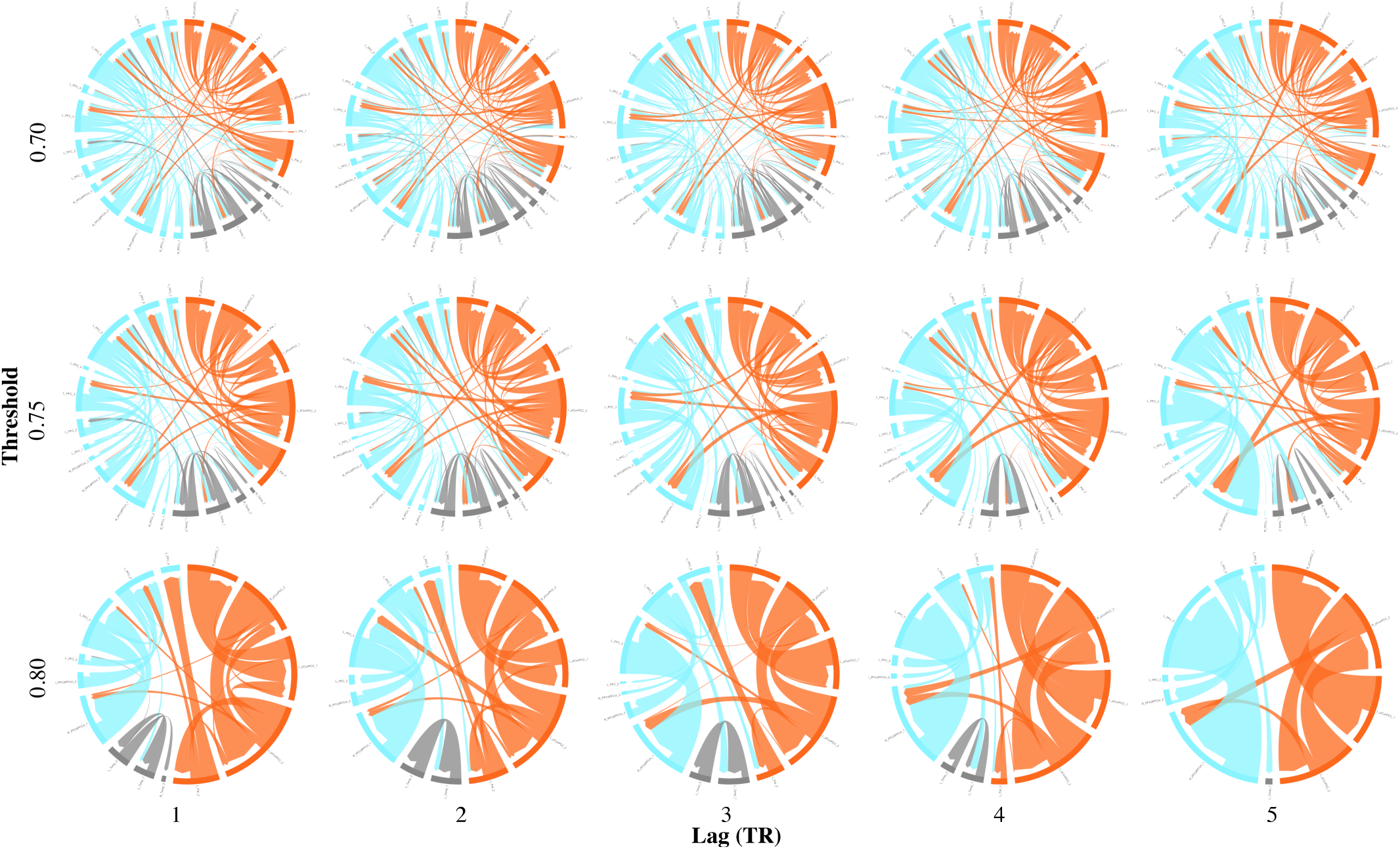
Effective connectivity within the DMN across lags and consistency thresholds. Reconstructed effective connectivity within the DMN for different negative lags ranging from 1 to 5 TRs (left to right) and different consistency thresholds from 0.7 to 0.8 (top to bottom). As shown in the main text, posterior-to-anterior asymmetric influences were consistent across hyper-parameters chosen in the analyses for reasonable thresholds and lags. Note how the number of regions with incoming or outgoing connections diminishes for long TRs and high thresholds.

## Footnotes

1 We used the statsmodels Python library 0.14.0.

